# Multi-Omics Analysis Reveals the Attenuation of the Interferon Pathway as a Driver of Chemo-Refractory Ovarian Cancer

**DOI:** 10.1101/2024.03.28.587131

**Authors:** Daria Afenteva, Rong Yu, Anna Rajavuori, Marina Salvadores, Inga-Maria Launonen, Kari Lavikka, Kaiyang Zhang, Giovanni Marchi, Sanaz Jamalzadeh, Veli-Matti Isoviita, Yilin Li, Giulia Micoli, Erdogan Pekcan Erkan, Matias M. Falco, Daniela Ungureanu, Alexandra Lahtinen, Jaana Oikkonen, Sakari Hietanen, Anna Vähärautio, Inderpreet Sur, Anni Virtanen, Anniina Färkkilä, Johanna Hynninen, Taru A. Muranen, Jussi Taipale, Sampsa Hautaniemi

## Abstract

Ovarian high-grade serous carcinoma (HGSC) represents the deadliest gynecological malignancy, with 10-15% of patients exhibiting primary resistance to first-line chemotherapy. These primarily chemo-refractory patients have particularly poor survival outcomes, emphasizing the urgent need for developing predictive biomarkers and novel therapeutic approaches. Here, we show that interferon type I (IFN-I) pathway activity in cancer cells is a crucial determinant of chemotherapy response in HGSC. Through a comprehensive multi-omics analysis within the DECIDER observational trial (ClinicalTrials.gov identifier NCT04846933) cohort, we identified that chemo-refractory HGSC is characterized by diminished IFN-I and enhanced hypoxia pathway activities. Importantly, IFN-I pathway activity was independently prognostic for patient survival, highlighting its potential as a biomarker. Our results elucidate the heterogeneity of treatment response at the molecular level and suggest that augmentation of IFN-I response could enhance chemosensitivity in refractory cases. This study underscores the potential of the IFN-I pathway as a therapeutic target and advocates for the initiation of clinical trials testing external modulators of the IFN-I response, promising a significant stride forward in the treatment of refractory HGSC.

---

Chemotherapy resistance is the leading cause of cancer-related deaths, representing the most critical unresolved challenge in oncology. The issue of chemotherapy resistance is particularly acute in ovarian high-grade serous carcinoma (HGSC), which is the deadliest gynecological malignancy that accounts for over 200,000 deaths per year worldwide^1,2^. HGSC is typically diagnosed at an advanced stage when dissemination from ovaries and fallopian tubes, which are considered the site-of-origin, into the peritoneal cavity has already occurred^3–5^, which hinders the effectiveness of treatments. Genomically, HGSC is a copy-number-driven cancer^6^ that is characterized by an almost 100% *TP53* mutation rate^7^ and high patient- and tissue-specific heterogeneity^8^, which further make HGSC challenging to treat, leading to < 40% five-year overall survival^9^.

The standard-of-care for HGSC is cytoreductive surgery followed by platinum-taxane chemotherapy and possible maintenance therapy with anti-angiogenesis or poly (ADP-ribose) polymerase (PARP) inhibitor^10^. The PARP inhibitors have significantly improved survival rates of patients with dysfunctional *BRCA1* or *BRCA2* and, more generally, patients with homologous recombination-deficient disease^11^. Patients who have a low likelihood for optimal surgical cytoreduction at diagnosis (40-50% of all patients with HGSC^12^) are referred to neoadjuvant therapy (NACT), which consists of 3-4 cycles of platinum-taxane chemotherapy followed by cytoreductive surgery, adjuvant chemotherapy, and possible maintenance therapy^10^. Notably, 10-15% of patients respond inconspicuously or not at all to NACT and cannot be operated. These primarily chemo-refractory patients have the worst prognosis of already poor prognosis HGSC and are in dire need of effective treatment options^13^.

To address this unmet clinical need, we present herein a unique prospective subcohort of 31 patients with HGSC who are primarily chemo-refractory and belong to the DECIDER trial^8^. We employed multi-omics data from these patients, including whole-genome sequencing (WGS), bulk and single-cell RNA-seq, and spatial data from highly multiplexed images to discover and validate molecular drivers of primarily chemo-refractory disease. Our results show that the platinum response is associated with baseline interferon type I (IFN-I) pathway activity, for the first time opening the avenue for personalized modification of the primary chemotherapy for patients with primarily chemo-refractory HGSC.

## Results

### Patient characteristics

We established a subcohort of 31 NACT-treated patients with chemo-refractory HGSC from the prospective, longitudinal, multi-region, observational DECIDER trial (Multi-Layer Data to Improve Diagnosis, Predict Therapy Resistance and Suggest Targeted Therapies in HGSOC; ClinicalTrials.gov identifier NCT04846933) (Fig. 1a,b, Extended Data Fig. 1 and Methods). The chemo-refractory disease was defined as a stable or progressive disease after primary therapy according to the RECIST 1.1 criteria^14^. Patients who did not receive at least two cycles of chemotherapy were excluded.

**Figure 1.**
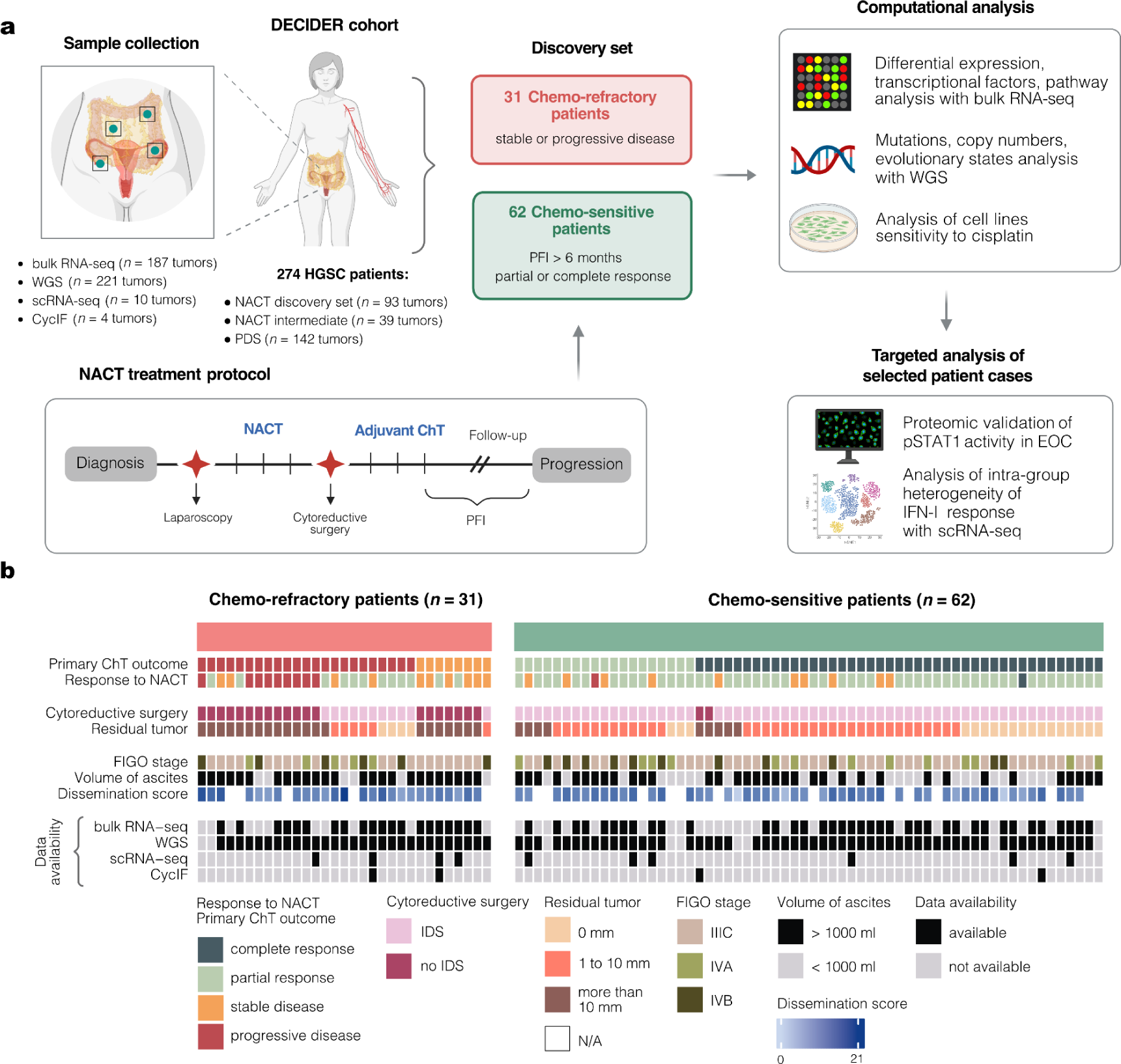
Study design and clinical characteristics of chemo-refractory and chemo-sensitive HGSC patients from the DECIDER cohort. **a**, Schematic representation of sampling procedure and study design. DECIDER cohort comprised patients treated with PDS and NACT, including chemo-refractory (*n* = 31), chemo-sensitive (*n* = 62), and intermediate patients (*n* = 39). **b**, Clinical characteristics of chemo-refractory (left) and chemo-sensitive (right) patients included in the discovery set and data availability. RNA-seq, RNA sequencing; WGS, whole-genome sequencing; scRNA-seq, single-cell RNA sequencing; CycIF, cyclic immunofluorescence; HGSC, ovarian high-grade serous carcinoma; NACT, neoadjuvant chemotherapy; PDS, primary debulking surgery; PFI, platinum-free interval; ChT, chemotherapy; pSTAT1, phospho-STAT1; IFN-I, interferon type I; FIGO, International Federation of Gynecology and Obstetrics; IDS, interval debulking surgery.

For comparison analyses, we selected 62 chemo-sensitive patients with similar baseline clinical characteristics, including the same treatment strategy but at least partial response to primary therapy and platinum-free interval (PFI), which is calculated from the last dose of platinum to first disease progression, exceeding six months (Fig. 1a,b, Extended Data Fig. 1 and Methods). The patient characteristics are shown in Supplementary Table 1. The refractory patients had a higher disease burden at the time of diagnosis based on higher dissemination score^15^ (*P* = 0.026) and more frequently detected large volume of ascites (*P* = 0.024) (Fig. 1b, Supplementary Table 1).

Diagnostic biopsies were available for 84 patients in the discovery set (Fig. 1b). Bulk RNA sequencing (RNA-seq) was performed on fresh frozen biopsies from 20 refractory and 38 chemo-sensitive tumors, a subset of which were selected for single-cell RNA-seq (*n* = 10) (Extended Data Fig. 1). Whole-genome sequencing (WGS) data from tumor and germline reference samples was available from 21 chemo-refractory and 41 chemo-sensitive patients. Cyclic immunofluorescence (CycIF) was performed on four patients (Extended Data Fig. 1).

### Genomic landscape of chemo-refractory and chemo-sensitive tumors

We compared mutations, copy numbers, structural variants, and mutational signatures using WGS data from treatment-naive chemo-refractory and chemo-sensitive tumors. All patients had dysfunctional *TP53*, and there were no significant differences in mutations of homologous recombination repair genes, foldback inversions, and homologous recombination deficiency (HRD)-associated somatic mutational signatures SBS3 and ID6 (Fig. 2a, Supplementary Table 1). Germline mutations of *BRCA1/2* (*n* = 3) and *RAD51C/D* (*n* = 2) were detected only in the chemo-sensitive patients, whereas a reversion mutation of *BRCA1* was detected in one chemo-refractory patient.

**Figure 2.**
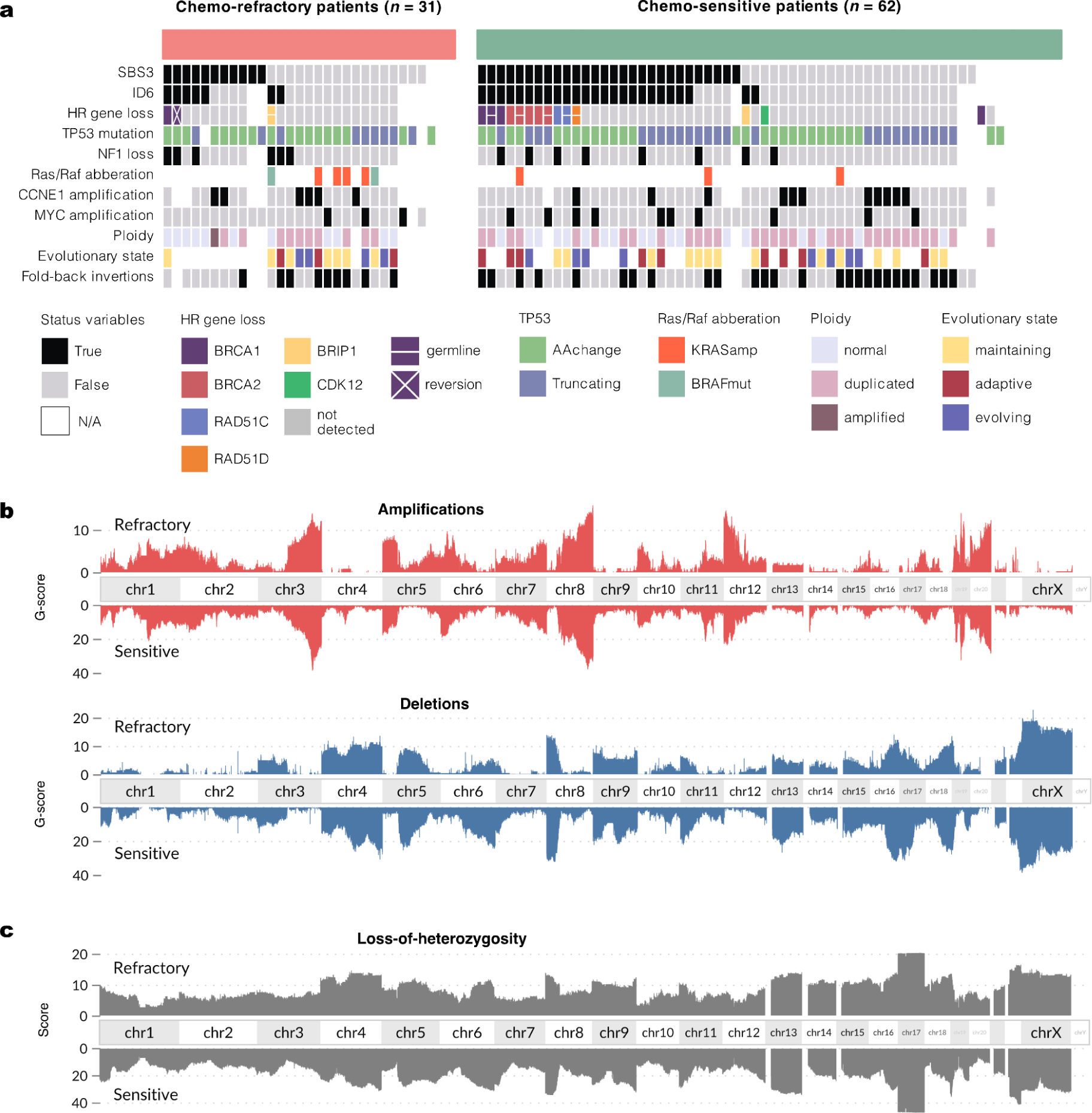
Genomic landscape of chemo-refractory and chemo-sensitive HGSC. **a**, Summary of genomic oncogenic events for the chemo-refractory (*n* = 31) and chemo-sensitive (*n* = 62) patient groups. **b**, Amplifications (red) and deletions (blue) across all chromosomes in chemo-refractory patients, with genomic alterations plotted relative to a reference genome of chemo-sensitive patients, revealing similar copy-number landscapes. **c**, Loss-of-heterozygosity across the genome of chemo-refractory and chemo-sensitive patients. The G-score was calculated as the total magnitude of aberrations (logR) across the genome^68^.

Established oncogenic driver aberrations, such as amplification of *KRAS*, *MYC*, or *CCNE1* and loss of *NF1,* were equally rare in both patient groups (Fig. 2a). Oncogenic *BRAF* class III kinase-impaired mutations were identified in two refractory patients, while none were found in sensitive patients. Notably, within the entire DECIDER cohort (*n* = 221 patients with WGS data), there were two additional patients with *BRAF* class III mutations: one patient whose primary NACT was canceled after one cycle due to lack of response and another patient with poor survival treated with primary debulking surgery (PDS), thus not included in the discovery analysis. Whole genome duplications were detected in > 50% of the samples with no marked distinction between the two groups (Fig. 2a). Additionally, the copy number profiles for gains, losses, and allelic imbalance were strikingly similar between the groups (Fig. 2b). The loss-of-heterozygosity (LOH) of the entire length of chromosome 17 was observed in all cancer samples (Fig. 2c). Tumor evolutionary states describing subclonal and site-associated heterogeneity were uniformly represented in the groups, as shown in the interactive GenomeSpy visualization of the genomic landscape of all samples^8,16^ (https://csbi.ltdk.helsinki.fi/p/chemorefractory/) (Fig. 2a-c).

### Low-interferon, high-hypoxia cell state is associated with chemo-refractoriness and prognosis

To identify processes that drive chemo-refractory phenotype, we first decomposed bulk RNA-seq data to cancer, immune, and stromal components with PRISM^17^ and performed a Differential Expression Analysis (DEA) for the cancer component of chemo-refractory and chemo-sensitive tumors (Methods, Extended Data Fig. 2a-d). Leveraging gene-level statistics from DEA, we conducted pathway activity inference using PROGENy^18^, evaluating enrichment scores for 14 pathways, and identified the perturbed transcriptional factors from transcriptomic data using CollecTRI^19^ (Methods, Extended Data Fig. 2a).

The largest activity difference was observed in the JAK-STAT pathway (*P* < 1e-16), primarily due to the reduced expression of more than 77% of its positive targets, such as *CXCL11*, *CMPK2*, *ISG15* (Fig. 3a,c). The most active pathway in chemo-refractory samples in comparison to chemo-sensitive was the hypoxia pathway (*P* = 3.3e-6) (Fig. 3a), regulated by *HIF1A*-dependent transcription (*P* = 1.6e-4) (Fig. 3b, Extended Data Fig. 2e). Interestingly, we observed that low JAK-STAT activity and high hypoxia were inversely related phenotypes (Spearman correlation coefficient *ρ* = −0.59, *P* = 6.1e-9) (Fig. 3d).

**Figure 3.**
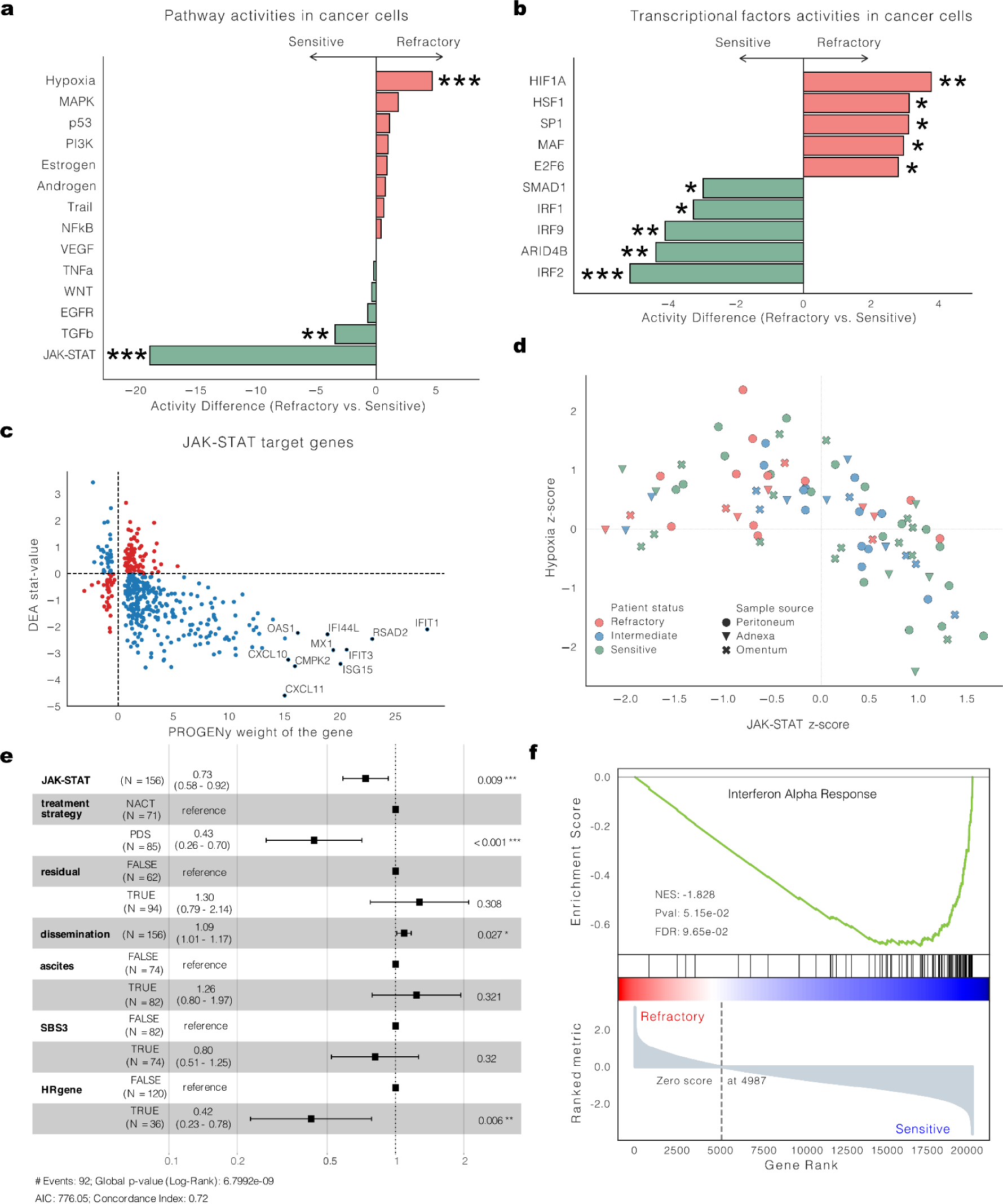
Pathway and transcriptional factors associated with chemo-refractoriness. **a**, Pathway activity differences between chemo-sensitive and chemo-refractory HGSC patients calculated with PROGENy revealed significant (two-tailed Student’s *t*-test < 0.05) down-regulation of JAK-STAT and TGFβ pathways, along with up-regulation of the hypoxia pathway in refractory patients. **b**, Differential activity of transcription factors between chemo-sensitive and chemo-refractory HGSC patients calculated with CollecTRI, highlighting the lower activity of interferon regulatory factors *IRF2*, *IRF9*, *IRF1,* and higher *HIF1A* activity in refractory patients. Top 5 significantly (two-tailed Student’s *t*-test < 0.05) up- and down-regulated transcriptional factors are presented. **c**, Scatter plot depicting the weight of the JAK-STAT target genes (*x*-axis) and their stat-value derived from Differential Expression Analysis (DEA) analysis (*y*-axis) of chemo-refractory versus chemo-sensitive tumors. **d**, Correlation plot of hypoxia and JAK-STAT z-scores across samples from chemo-refractory (*n* = 20), intermediate (*n* = 24), and chemo-sensitive (*n* = 38) patients from the DECIDER cohort, indicating an inverse relationship. Data points are color-coded by patient status and symbol-coded by tissue origin. One representative treatment-naive sample from solid metastatic or intra-abdominal tissues with the lowest JAK-STAT score was taken per patient (Supplementary Table 3 and Methods). **e**, Multivariable Cox proportional hazards model showing the prognostic significance of the JAK-STAT score for overall survival (*n* = 156 patients) in NACT and PDS patients from the DECIDER cohort. One representative treatment-naive sample from solid metastatic or intra-abdominal tissues with the lowest JAK-STAT score was taken per patient (Methods). The whiskers represent the 95% CI. The residual was classified as TRUE when the residual tumor after cytoreductive surgery was more than 0 mm. HRgene indicates the presence of a mutation in the homologous recombination deficiency (HRD)-related genes. SBS3 indicates the HRD status of a patient according to the SBS3 mutational signature. **f**, GSEA indicating significant enrichment of the Interferon Alpha Response pathway, with significantly lower activity in chemo-refractory patients, highlighted by the negative normalized enrichment score (NES). The Benjamini–Hochberg (BH) procedure was used to adjust the two-sided *P* values for multiple testing for GSEA. \**P* < 1e-2, \*\**P* < 1e-3, \*\*\**P* < 1e-5 (two-tailed Student’s *t*-test).

To test whether the JAK-STAT pathway has an association with patient survival, we calculated PROGENy scores for the JAK-STAT pathway for all patients with bulk RNA-seq data from the DECIDER cohort (*n* = 156) and fitted the multivariable Cox proportional hazards model (Methods, Fig. 3e). The JAK-STAT score was prognostic for overall survival (HR = 0.73, 95% CI = 0.58-0.92, *P* = 0.009). For NACT patients stratified into three groups by JAK-STAT activity scores, Kaplan-Meier survival analysis (Extended Data Fig. 2g) revealed a survival advantage in the high-activity group over the low-activity group (*P* = 0.007).

To identify the perturbed axis of the JAK-STAT pathway in chemo-refractory tumors, we analyzed the activity of transcriptional factors, which revealed that interferon regulatory factors *IRF2*, *IRF9*, and *IRF1* had lower activity in refractory compared to the chemo-sensitive tumors (Fig. 3b). The reduction in *IRF9* activity, a pivotal component of the interferon-stimulated gene factor-3 (ISGF3) complex alongside *STAT1* and *STAT2*, directly implicates suppressed IFN-I response. The attenuation of this pathway in the chemo-refractory patients was further evidenced by decreased expression levels of the *IRF9*-regulated genes *CXCL10, IFIT3, OAS1, IFIT1,* and *STAT1* (Extended Data Fig. 2f). Furthermore, the Gene Set Enrichment Analysis^20^ (GSEA) of the Molecular Signatures Database^21^ (MSigDB) Hallmark collection indicated the Interferon Alpha Response pathway as the most enriched (Benjamini–Hochberg (BH)-adjusted *P* = 0.097, NES = −1.828) in the genes ranked by the *t*-statistic (Fig. 3f). These analyses demonstrate that IFN-I response is suppressed in the chemo-refractory HGSC patients at the time of diagnosis.

### Genomic perturbations do not explain reduced IFN-I activity in chemo-refractory patients

To investigate whether genetic aberrations are responsible for the attenuated IFN-I signaling in chemo-refractory patients, we conducted an in-depth analysis of somatic aberrations of the genes implicated in the IFN-I response cascade (Extended Data Fig. 3a,c). The genomic landscape was devoid of loss-of-function mutations, except for a deletion in the IFN-alpha/epsilon and *CDKN2A/B* locus of one patient (Extended Data Fig. 3b). Our results, or lack thereof, suggest that genomic alterations are unlikely to be the primary driver of the diminished IFN-I response characterizing chemo-refractory patients.

### Single-cell RNA-seq and spatial protein profiling of the IFN-I activity confirm its association with chemo-refractoriness

We then characterized the IFN-I activity and heterogeneity using single-cell transcriptomic and spatial data from highly multiplexed images. Single-cell RNA-sequencing (scRNA-seq) data were acquired from four refractory and six chemo-sensitive patients (Fig. 4a). Employing a tiered clustering approach^22^, we catalogued 16,682 high-quality cells, including malignant (*n* = 4,577), stromal (*n* = 2,143), and immune cells (*n* = 10,062) (Fig. 4b,c).

**Figure 4.**
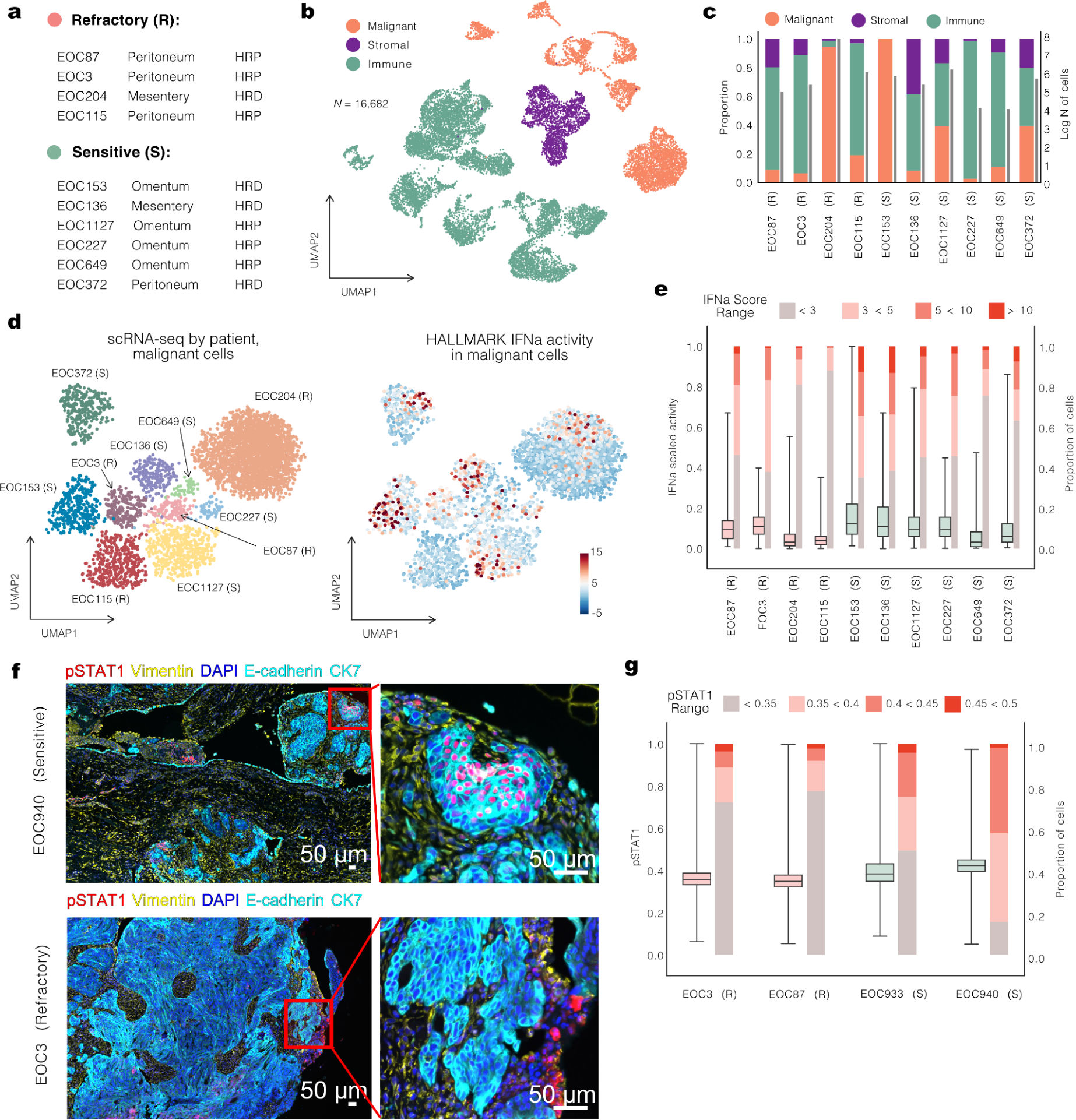
Single-cell RNA-seq and CycIF analyses of selected patients reveal the heterogeneity of IFN-I activity in chemo-sensitive patients. **a**, Overview of single-cell RNA sequencing (scRNA-seq) data from solid metastatic tissues from four refractory (R) and six chemo-sensitive (S) HGSC patients. Homologous recombination deficiency (HRD) status was defined by the SBS3 mutational signature. **b**, UMAP of 16,682 scRNA-seq profiles from ten patients colored by cell type. **c**, Proportions of different cell types (left axis) and logarithmic number of cells (right axis) per tumor. **d**, UMAP of 4,577 malignant cells colored by the patient (left) and by the level of IFN-I activity calculated with DecoupleR (right). **e**, Boxplots illustrating normalized IFN-I pathway activity scores in individual cells within each tumor, showing a broader spread of activity in chemo-sensitive patients. Bar plots show the proportion of cells with IFN-I activity scores within a specific range for every tumor. **f**, Tissue cyclic multiplex immunofluorescence (t-CycIF) imaging of FFPE tumor samples from the omentum and peritoneum of chemo-refractory (EOC3) and chemo-sensitive (EOC940) patients, respectively, with pSTAT1 expression indicating IFN-I activity. Scale bar, 50 μm. **g**, Boxplots with pSTAT1 expression in malignant cells across chemo-refractory (EOC3, EOC87) and chemo-sensitive (EOC933, EOC940) patient samples. Bar plots show the proportion of cells with pSTAT1 expression within a specific range for every tumor. Box plots are presented as the range (whiskers) with the bounds of the box extending to the first and the third quantiles with a line at the median and whiskers extending to the maximal and minimal data points (Fig. 4e,g). HRP, homologous recombination proficiency; pSTAT1, phospho-STAT1.

To address the intra-patient variability of the IFN-I activity, we computed the IFN-I pathway score for each single cell sample and observed distinct patterns of its activation across various cell types. The IFN-I pathway exhibited elevated activity in malignant cells of the chemo-sensitive patients compared to the refractory patients, corroborating the bulk RNA-seq results (Fig. 4d). Furthermore, we found that the cancer cells from chemo-sensitive samples encompassed a more diverse spectrum of the pathway activity, which manifested in a higher variance of the activity scores with the presence of a subset of cells in the extended rightward tail in the distribution (Fig. 4e).

We next performed a single-cell analysis of formalin-fixed paraffin-embedded (FFPE) tumor samples from omental and peritoneal regions of two refractory and two sensitive patients using multiplexed cyclic immunofluorescence (t-CycIF) imaging^23^. We annotated approximately 1.4 million cells into cancer, immune, and stromal cells using TRIBUS^24^. We assessed the state of IFN-I pathway activity by quantifying the expression levels of its indicator phospho-STAT1 (pSTAT1). Spatially, the pSTAT1-positive cancer cells formed clusters in the chemo-sensitive tumors, whereas similar colocalization was not observed in the chemo-refractory tumors, where the great majority of cancer cells were negative for pSTAT1 (Fig. 4f,g). In line with the results from scRNA-seq, t-CycIF analysis confirmed an elevated IFN-I pathway activity in the cancer compartment of chemo-sensitive tumors with the presence of malignant cells distinguished by highly active IFN-I response (Fig. 4g).

### Platinum response of ovarian cancer cell lines is associated with IFN-I signaling

We then investigated the role of IFN-I signaling in platinum resistance in Kuramochi and COV362 cell lines (Supplementary Table 2), which are genomically similar to HGSC tumors^25^. We performed scRNA-seq combined with cell hashing and determined transcriptomic profiles of the HGSC cells exposed to different concentrations of platinum (10%, 50%, and 200% of IC50) for different time intervals (24h/12h) (Fig. 5a). We first calculated the effect difference between platinum-treated and control cells by extracting the platinum gene expression signature from scRNA-seq data using Cohen’s *d* (Methods, Extended Data Fig. 4a). The extracted platinum signatures showed a consistent correlation across the replicates (*n* = 5 for each cell line) (Extended Data Fig. 4a). A platinum sensitivity score (CisSenScore, Methods) for each cell was calculated and based on it, the platinum-treated cells were classified into ‘more-sensitive’ and ‘less-sensitive’ groups (Fig. 5b, Extended Data Fig. 4b).

**Figure 5.**
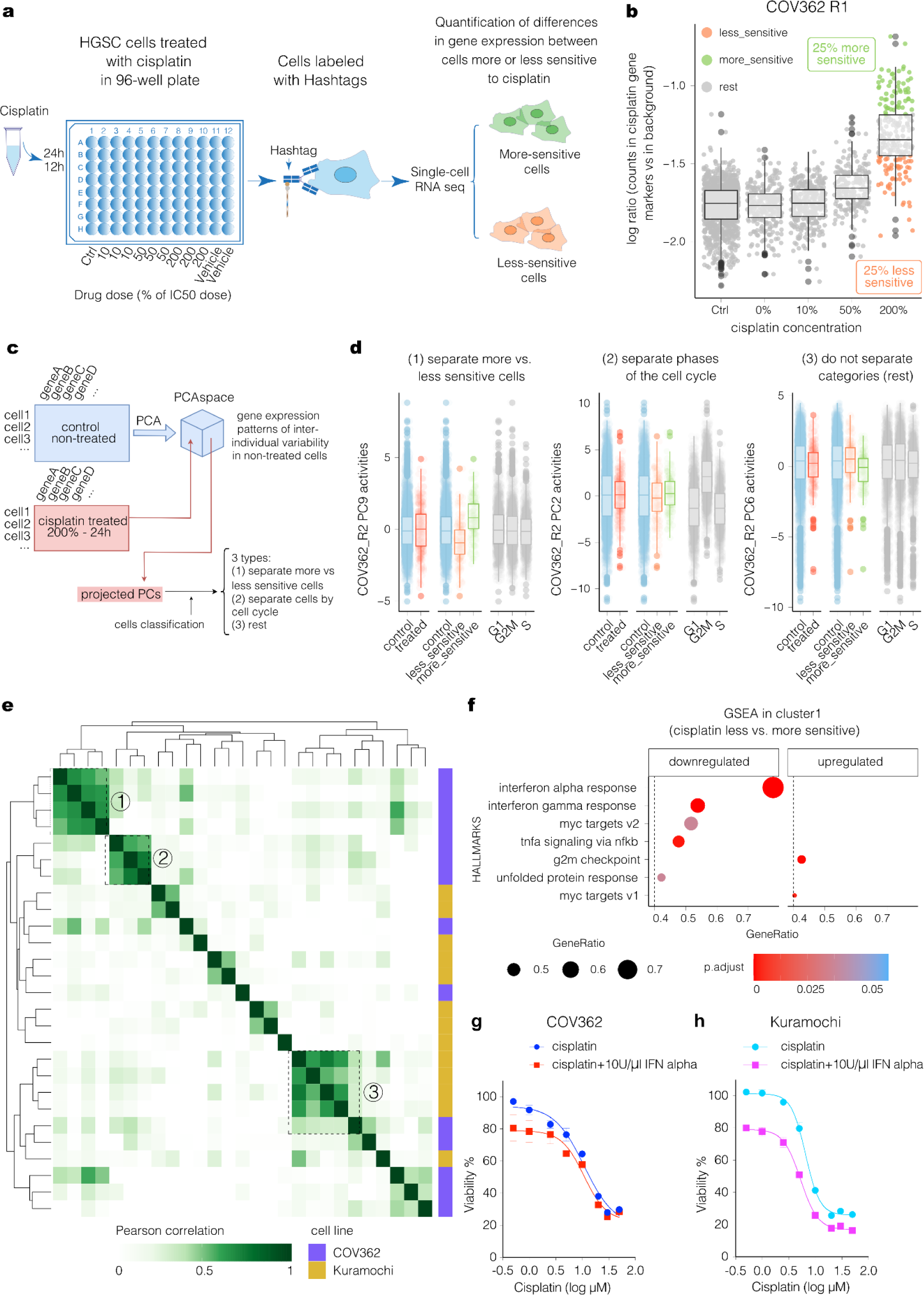
Single-cell RNA-seq in HGSC cell lines shows that IFN-I response determines platinum-based drug resistance. **a**, Workflow of the single-cell RNA-seq combined with cell hashing in cell lines. **b**, Based on the CisSenScore, the treated cells were separated into ‘more-sensitive’ (top 25%, green) and ‘less-sensitive’ (bottom 25%, orange) in COV362 cells. R1 refers to one of the experiment replicates. **c**, Schematic of the data analysis pipeline. The non-treated cells underwent PCA to extract gene expression patterns of inter-individual variability, and the treated cells were projected into the PC space to be classified into three types based on the CisSenScore. **d**, Three types of PCs in treated cells of COV362_R2 experiment: (1) able to separate ‘more-sensitive’ vs ‘less-sensitive’ cells; (2) able to separate different phases of the cell cycle; (3) not able to separate. **e**, Heatmap depicting pairwise similarities between gene expression patterns of inter-individual variability identified in non-treated cells across replicates. Hierarchical clustering identified 3 clusters indicated by dashed boxes and numbers. **f**, GSEA scores for hallmark gene sets (with p.adjust < 0.05 and GeneRatio > 0.4) were calculated based on the gene expression signature in cluster1 of the ‘less-sensitive’ cells compared with the ‘more-sensitive’ cells. Results from cluster 1 are shown. **g**, Cell viability of COV362 cells treated with different concentrations of cisplatin-only or cisplatin combined with 10U/μl IFN-alpha. **h**, Cell viability of Kuramochi cells treated with different concentrations of cisplatin-only or cisplatin combined with 10U/μl IFN-alpha. Box plots are presented as the range (whiskers) with the bounds of the box extending to the first and the third quantiles with a line at the median and whiskers extending to the farthest data point lying within 1.5x the interquartile range from the box.

To identify cellular mechanisms influencing platinum response in an unbiased fashion, we conducted principal component analysis (PCA) on scRNA-seq data from untreated cells, followed by projection of the expression data from treated cells onto the principal component (PC) state established by control cells’ expressions. This allowed the identification of PCs that capture the variation related to the platinum response. We selected the PCs with high variance effect size between ‘more-sensitive’ and ‘less-sensitive’ cells (not separating cell cycle phases; Methods) (Fig. 5c,d). The resulting 27 PCs were clustered using hierarchical clustering, yielding three clusters (Fig. 5e). Analysis of the three clusters using GSEA^20^ showed that clusters 1 and 2 in the ‘less-sensitive’ cells had a significantly down-regulated activity of the Interferon Alpha Response pathway, corroborating the IFN-I activity reduction in patients with chemo-refractory HGSC (Fig. 5f, Extended Data Fig. 4c-e).

Next, we investigated the effect of IFN-alpha and platinum combination treatment on cell viability. We tested six IFN-alpha concentrations for the combination treatment and selected the 10U/μl concentration (Extended Data Fig. 4g). The combined treatment significantly reduced the cell viability compared with platinum-only treatment and showed additive effects of IFN-alpha and platinum in both COV362 and Kuramochi cell lines (Fig. 5g,h). Taken together, these results indicate that the IFN-I signaling state of HGSC cells is intrinsically variable and that high levels of IFN-I signaling cell-autonomously increase the chemosensitivity of the cells to platinum.

## Discussion

Molecular processes driving chemo-refractory disease remain elusive, and currently, no biomarkers or effective therapy options exist for patients with HGSC who do not respond to first-line chemotherapy. To address this clinically urgent but unmet problem, we established multi-omics data from clinically curated chemo-refractory and corresponding chemo-sensitive patients with HGSC within the prospective, observational DECIDER clinical trial. All samples harbored *TP53* mutation as expected, and an experienced gynecopathologist verified that samples with atypical genomes were indeed serous carcinomas.

Herein, we demonstrate that the basal activity of the IFN-I in cancer cells is associated with response to chemotherapy. Moreover, our findings indicate substantial heterogeneity in IFN-I activity in the treatment-naïve HGSC tumors, suggesting that high variability in this pathway activity contributes to inherent heterogeneity in treatment response. The scRNA-seq and CycIF data, as well as the functional cell line experiments, replicated the finding of the baseline heterogeneity in susceptibility to platinum, with cells being more sensitive to cisplatin having elevated activity of IFN-I compared with more resistant cells. Combination treatment experiments with cell lines also suggest that the effect of IFN-I response is primarily cell-autonomous, appearing less dependent on the tumor microenvironment. Importantly, we showed that IFN-I activity in cancer cells is an independent prognostic marker of poor response to the first-line therapy in HGSC.

Recent work on molecular characterization of adaptation and acute response to PARP inhibitors suggested that IFN signaling upregulation is an early response^26^. In the highly adapted states, silenced IFN signaling is accompanied by elevated *HIF1A*-signaling, contributing to the management of oxidative stress. Based on our findings, the chemo-refractory HGSC phenotype resembles these far-adapted cell states, with low baseline IFN-I activity and high hypoxia-related signaling.

Our results suggest that patients with chemo-refractory HGSC may benefit from inducing IFN-I activity to enhance their responsiveness to chemotherapy. This treatment suggestion is supported by a phase II study that combined cisplatin and alpha-2 interferon in non-small cell lung cancer, with a 30% response rate and acceptable toxicity^27^. Indeed, several approaches to increase IFN-I activity have been published recently, such as targeting macrophages^28^ or IFN-epsilon^29^. As chemo-refractory patients currently do not have efficient treatment options, a clinical trial to test the efficacy of IFN-I modulation and platinum-based chemotherapy is warranted.

Despite thorough genetic analysis of our unique dataset, we did not find genetic drivers behind chemo-refractory phenotype or lack of IFN-I activity, hinting at non-genomic mechanisms behind chemotherapy resistance. Unlike Chowdhury and colleagues^30^, we did not find the LOH of chromosome 17 to be enriched in patients with chemo-refractory HGSC. However, we replicated their finding that *BRAF*-mutated tumors were exclusively found in the refractory group. In the full DECIDER cohort, all four patients with *BRAF* mutations had poor survival and Class III mutations. The Class III *BRAF* mutations impair the kinase domain but retain partially some RAS-dependent activity. Cancers with such mutations have been demonstrated to be sensitive to MEK, ERK, or RAS inhibition^31^. While the prevalence of *BRAF* mutations in HGSC is ∼2.6%^31^, accumulating evidence indicates that the patients harboring *BRAF* mutations do not respond to first-line chemotherapy. As a significant benefit from targeted therapy with MAPK inhibitors has been demonstrated for cancers with Class III *BRAF* mutations, patients with primary chemo-refractory HGSC should be tested for *BRAF* mutations.

We recognize the limitations of this study. A mechanistic explanation for the lack of IFN-I activity in chemo-refractory patients remains to be elucidated. While our data suggest that driving mechanisms for IFN-I are likely non-genomic, functional experiments with preclinical models that address tumor microenvironment are warranted. Furthermore, we acknowledge the need for validation of the results in independent patient cohorts.

In conclusion, we established IFN-I pathway activity in cancer cells as a critical determinant of the response to chemotherapy, offering a prognostic biomarker and the first therapeutic target to combat chemo-refractoriness in HGSC. By providing a detailed examination of the underrepresented patient population in ovarian cancer research, our results advocate for the urgent initiation of clinical trials to evaluate IFN-I modulation strategies, potentially transforming the treatment landscape for patients with chemo-refractory HGSC.

## Methods

### Study participants

The DECIDER study (Multi-Layer Data to Improve Diagnosis, Predict Therapy Resistance and Suggest Targeted Therapies in HGSOC; ClinicalTrials.gov identifier NCT04846933) fosters ongoing prospective recruitment of patients diagnosed with HGSC at the Turku University Hospital, which started in 2010. Clinical data on patient characteristics were collected from hospital records. For these analyses, we included patients diagnosed before 2022-03-31. The primary treatment strategy of all patients adhered to the ESMO guidelines^10^, including either primary debulking surgery (PDS) followed by platinum-based adjuvant chemotherapy, or neoadjuvant chemotherapy (NACT), interval debulking surgery (IDS), and adjuvant chemotherapy. The chemo-refractory study group included patients from the NACT-treated arm and was defined by their outcome from primary therapy with either stable or progressive disease, according to the RECIST version 1.1 criteria^14^. The platinum-free interval (PFI) of these patients was at highest 45 days, adhering to the ESMO definition of chemo-refractory^13^ HGSC with a slight concession for delayed progression detection. We excluded eleven patients from the analyses, most of whom did not receive adequate NACT treatment, comprising at least two cycles of chemotherapy due to pre-existing medical conditions or severe side effects. The reference group consisted of chemo-sensitive patients with similar baseline characteristics and treatment strategy but with either partial or complete response to primary therapy and a PFI of more than six months. This patient selection yielded 31 chemo-refractory and 62 chemo-sensitive cases. The full DECIDER cohort, including 39 additional NACT-treated patients with intermediate follow-up and 142 patients treated with PDS and adjuvant chemotherapy, was used to validate the findings from the discovery set (Supplementary Table 1).

### Sample preparation and selection

Tissue specimens from tumors were collected during diagnostic laparoscopy before treatment or palliative ascites removal and were subsequently subjected to pathological examination. Peripheral blood samples or buffy coat extracts were extracted in Auria Biobank for all patients for DNA extraction and genomic sequencing to serve as germline reference for identifying somatic genomic aberrations using Chemagic DNA Blood Kit Special (PerkinElmer Inc., USA) and Chemagic 360 instrument (PerkinElmer Inc., USA). For other samples, to extract DNA and RNA simultaneously, we utilized the Qiagen AllPrep kit (#80204). Slides of formalin-fixed paraffin-embedded (FFPE) tissue were stained with hematoxylin and eosin, scanned, and re-evaluated by a gynecopathologist (AVi). Tumor samples subjected to bulk and single-cell RNA-seq analyses originated from primary tubo-ovarian sites (ovaries and fallopian tubes) and solid metastatic sites (omentum, mesentery, and peritoneum). As many subsequent analyses required one sample per patient, we prioritized the samples from the solid metastatic sites. When multiple samples from solid metastatic sites were available for a patient, we selected the one with the highest abundance of EOC, according to PRISM^17^. To compare bulk RNA-seq data from chemo-refractory and chemo-sensitive tumors, we utilized samples from fresh frozen solid tumor biopsies listed in Supplementary Table 3.

### Whole-genome and RNA sequencing

Tissue samples were extracted from fresh frozen tissue, and those with sufficient DNA/RNA content were sent to BGI (BGI Europe A/S, Denmark) or Novogene (Novogene Europe, UK) for library preparation and nucleotide sequencing. Whole-genome sequencing (WGS) was performed with either DNBSEQ (BGISEQ-500 or MGISEQ-2000, MGI Tech Co., Ltd., China), HiSeq X Ten (Illumina, USA), or NovaSeq 6000 (Illumina, USA) as 100bp or 150bp paired-end sequencing with a median coverage of 47x. RNA sequencing was performed using DNBSEQ, HiSeq X Ten, HiSeq 4000, or NovaSeq 6000 as 100bp or 150bp paired-end sequencing.

### Bulk RNA-seq analysis

#### RNA-seq preprocessing

Bulk RNA sequencing (RNA-seq) reads were processed using the SePIA pipeline^32^ within Anduril2^33^, as was previously described in detail^17^. Trimmomatic^34^ v0.33 was used to trim low-quality bases. Trimmed reads were aligned to GRCh38.d1.vd1 with GENCODE v25 annotations via STAR aligner^35^ v2.5.2b, allowing up to 10 mismatches. Transcripts per million (TPM) and gene-level effective counts were quantified using eXpress^36^ v1.5.1-linux_x86_64. We applied the POIBM method for batch-effect correction to gene-level read counts^37^.

#### Bulk RNA-seq decomposition

We employed the latent statistical framework PRISM to extract the sample composition, scale factors, and cell-type-specific whole-transcriptome profiles adapted to each transcriptomic sample^17^. For the single-cell reference, we utilized single-cell RNA-seq data from HGSC, annotated for EOC, Fibroblasts, Immune, and Other, encompassing data from eight patients and various anatomical sites. Raw read counts were utilized as input for the model. This approach enabled us to accurately generate EOC-specific gene-level read counts for each bulk RNA sample.

#### Differential expression analysis

Prior to differential expression analysis (DEA), we refined the matrix of raw read counts by filtering out genes that were not adequately profiled. The criteria for retaining genes in the analysis were as follows: (1) the gene must have a minimum of 15 total reads across samples within each comparison group, and (2) the gene must have at least 10 counts in at least some samples within each group. This filtering process resulted in an input matrix comprising 15,730 genes across 58 samples. For the DEA, we utilized the Python implementation of the DESeq2 framework pydeseq2^38^ v0.3.4. This analysis aimed to identify genes differentially expressed between the chemo-refractory and chemo-sensitive tumors, with SBS3 and refractory/sensitive status as design factors in our model. We incorporated a unique approach for determining size factors, as follows: we utilized scale factors derived for each RNA sample through the PRISM framework multiplied by the EOC abundance and normalized by the geometric mean across all samples (https://github.com/XXX).

#### TF and pathway activity inference

To assess the pathway activity, we employed the Python implementation of the DecoupleR^39^ framework v1.5.0 and retrieved the PROGENy^18^ model weights with *decoupler.get_progeny(top=500)*. The multivariate linear model method *decoupler.run_mlm()* was used to infer the pathways enrichment scores. In addition to pathway activity scores, we also evaluated the activity scores of transcriptional factors (TF). For this purpose, we retrieved the CollecTRI^19^ gene regulatory network with *decoupler.get_collectri()*. For the inference of enrichment scores of TF, we utilized the univariate linear model method executed via *decoupler.run_ulm()*. We used the gene-level statistic incorporated in the stat-value of the DESeq2 output as input for this analysis.

#### GSEA

To infer functional enrichment scores, Gene Set Enrichment Analysis (GSEA) was conducted on the Trimmed Mean of M-values (TMM) normalized EOC expression profiles obtained using the Python package conorm v1.2.0. We utilized the hallmark gene sets from the Molecular Signatures Database^21^ (MSigDB) collection for this analysis. The GSEA was performed using the gseapy Python library v1.0.3, applying the following parameters: ranking method set to ‘t_test,’ a total of 1000 permutations for robust statistical assessment, and the inclusion criteria for gene sets were defined with a minimum of 15 genes and a maximum of 500 genes per set. We considered a pathway significantly enriched if Benjamini–Hochberg (BH)-adjusted *P* < 0.1.

#### Cell type abundance estimation

The Kassandra^40^ algorithm (tumor model) was used to evaluate the proportions of different cell types in the TME for each bulk RNA sample. TPM values obtained during the expression quantification phase served as the input data. When gene annotations were absent, zeroes were inputted to meet the algorithm’s input criteria.

### Single-cell RNA-seq analysis

Tumor samples were collected at the time of laparoscopy, dissociated into single-cell suspensions, and frozen for later scRNA-seq processing. The scRNA-seq libraries were prepared using the Chromium Single-Cell 3′ Reagent Kit v2.0 (10x Genomics) and were subsequently sequenced using the Illumina HiSeq 4000 at the Jussi Taipale Lab at the Karolinska Institutet, Sweden, as well as the HiSeq 2500 and NovaSeq 6000 at the Sequencing Unit of the Institute for Molecular Medicine Finland. Preprocessing of the data, comprising sample demultiplexing, alignment, barcode processing, UMI quantification, and cell type annotation, was performed as previously described^22^. To assess the IFN-I pathway activity, we employed the Python implementation of the DecoupleR^39^ framework v1.5.0 and retrieved the hallmark gene sets from MSigDB^21^ with *decoupler.get_resource(“MSigDB”)*. For the inference of enrichment scores of the IFN-I response pathway, we ran the Over Representation Analysis (ORA) executed via *decoupler.run_ora()*.

### WGS data analysis

#### WGS data preprocessing

The genomic DNA data processing was performed with the Anduril2 workflow platform^33^ and included quality control, alignment to the reference human genome, deduplication, cross-sample contamination estimation, and variant discovery. We used FastQC^41^ v0.11.4 and Trimmomatic^34^ v0.32 for sequenced DNA read quality control and trimming steps. Subsequently, the high-quality reads underwent alignment to the reference human genome GRCh38.d1.vd1 using BWA-MEM^42^ v0.7.12-r1039 with default parameters, subjected to deduplication with Picard v2.6 (https://broadinstitute.github.io/picard/) and base quality recalibration using the Genome Analysis Toolkit^43^ (GATK) v4.1.9.0. Additionally, cross-sample contamination estimation was conducted with GATK, setting the contamination estimation threshold at 3% for tumors, 5% for normals in Panel of Normals (PoN), and 10% for normals in germline variant calling.

#### Mutation calling

Somatic short variants were identified by collectively analyzing multiple tumor samples against a single-matched normal sample for each patient. This analysis was performed using GATK v4.1.9.0 Mutect2^44^, following established best practices^45^. The Finnish gnomAD^46^ v3.0 allele frequencies served as the germline resource for variant calling. We used a PoN generated with 181 normal samples from the DECIDER study and 99 TCGA normals^8^. Afterward, the GATK FilterMutectCalls tool was employed for variant filtration, retaining only those variants that successfully passed all applied filters. Variant allele frequencies (VAF) were computed by considering the read depths for both reference and alternate alleles as indicated in the AD field. We annotated the variants using the GATK VariantAnnotator, adding dbSNP^47^ 155 IDs. Additionally, an offline version of Combined Annotation Dependent Depletion^48^ (CADD) v1.6, complemented by an in-house solution, was employed to annotate variant call format files. The annotation process was further enriched using ANNOVAR^49^ 20191024, adding refGene^50^ data dated 2020-08-16, together with other annotations adapted for ANNOVAR, including ClinVar^51^ 20220816, customized annotations for COSMIC^52^ v96, and gnomAD v3.0 genomes. Germline variants were called as previously described^8^. The allele depths of variants at exons, untranslated regions, and splice sites were estimated by forced calling using GATK 4.1.9.0 Mutect2 in joint calling mode, as in somatic variant calling.

#### Tumor fraction estimation

Tumor fraction was estimated using a modified ASCAT^53^ algorithm that inputs copy-number segmentation from GATK^45^, as previously described^16^.

#### Copy number and breakpoint calling

Breakpoint and copy-number segmentation were determined by employing a combination of GRIDSS^54^ and the Hartwig Medical Foundation (HMF) toolkit. The pipeline was constructed on the Nextflow^55^ platform, utilizing default settings for the incorporated tools. Breakpoint calling was executed using GRIDSS^54^ v2.13.2, with the exclusion of regions found in both the ENCODE^56^ and in-house built DECIDER blacklists^8^. Subsequently, breakpoint filtering was accomplished through GRIPSS^57^ v2.0, leveraging a PoN originating from blood samples collected from DECIDER patients and a Dutch population. B allele frequency (BAF) calculation was conducted using the Amber tool v3.8, while heterozygous biallelic loci were identified as described in the preceding section. Cobalt v1.12 was employed to assess read depth normalized for GC content. The combined results from all these tools, in addition to somatic single nucleotide variants (SNVs), were then used to input data into PURPLE^57,58^ v3.7.2. PURPLE estimated copy-number segmentation profiles and conducted estimations for tumor purity and ploidy of the samples. Breakpoints derived from GRIDSS-PURPLE analysis were used for structural variants annotation with Linx^59^ v1.22. Among all events, simple and chained foldback inversion were extracted and used for the analysis.

#### Mutational signatures and signature events

Mutational signatures were fitted, and SBS3- and ID6-based homologous recombination deficiency (HRD) statuses were computed as previously described^60^ using COSMIC^61^ v3.3.1 reference signatures. A cancer was classified as positive for foldback inversions if at least one tissue sample was positive. A sample sequenced with NovaSeq was called positive if at least five foldback inversion events were detected, and a sample sequenced with BGISEQ or HiSeq was called positive if at least three foldback inversion events were detected to accommodate the platform-specific differences in detection sensitivity.

#### Curation of somatic aberrations

To define the genomic status of genes involved in the IFN-I signaling cascade, we curated function-affecting mutations and the dosage of intact gene copies and summarized the dosage perturbation over the whole signaling cascade. First, we collected short somatic mutations and annotated, weighing a mutation a loss-of-function if its consequence was protein truncation or if it was a missense predicted deleterious unanimously by polyphen2^62^ and SIFT^63^. Then, we retrieved all genomic breaks from the PURPLE-processed data and assessed their effect on the coding sequence. With locus copy number and tumor fraction in the sample from the processed data and with read counts supporting the aberration, we estimated the number of gene copies affected by the short or structural aberrations and the number of intact gene copies. Using the most common copy number across the cancer genome as a reference copy number, we calculated the dosage of intact gene copies as a ratio of the number of intact gene copies and the reference copy number. Gene dosage below or equal to 0.5 or above or equal to 2.0 was considered perturbed. Adapting the molecular distance introduced earlier^64^, a pathway perturbation score of cancer was calculated as a log2-scale sum of absolute perturbed gene dosages.

### Tissue cyclic multiplex immunofluorescence (t-CycIF)

Whole-slide HGSC FFPE samples were stained and scanned iteratively with validated antibodies (DAPI, E-Cadherin (CST, CAT 3199), CK7 (Abcam, CAT ab209601), Vimentin (CST, CAT 9855), pSTAT1 (CST, CAT 8183S)) using RareCyte CyteFinder scanner following the t-CycIF workflow^65^ as described^23^. BaSiC tool and the Ashlar algorithm were used for image correcting, stitching, and registration^66^. Single-cell nuclei segmentation was performed using probability maps created by UNET^67^, and these were dilated by 2px to obtain whole-cell segmentation masks. The mean fluorescence intensity for each cell was computed using in-house scripts and Python’s scikit-library to obtain a single-cell data table. SOM-based TRIBUS^24^ algorithm was used for cell type calling, and pSTAT1 expression was gated and rescaled (*pl.gate_finder* and *pp.rescale* using the scimap package in Python) to a value between 0 and 1 separately for each image so that values above 0.5 identify cells expressing the marker as described^23^.

### Statistical analyses

Statistical tests for patient and cancer characteristics were performed with the Fisher exact test for categorical variables and the Mann-Whitney *U*-test for ordinal variables. Statistical tests for transcriptional factors and pathway activities were performed using the Student’s *t*-test. All *P* values were two-sided. The BH procedure was used to adjust the two-sided *P* values for multiple hypothesis testing when appropriate. The strength of correlations was measured using the Spearman (*ρ*) correlation coefficient and the probability of observing a correlation with the corresponding *P* values. The threshold for significance was set up as less or equal to *P* = 0.05 in all analyses unless specified otherwise.

Patient survival was analyzed using R library survival, with a diagnosis of HGSC as a starting point of time at risk and death from HGSC as an endpoint. The follow-up was cut at 2023-01-31 and patients were censored at that time, if alive. One patient was censored at a time of non-HGSC-related death. Hazard ratios were estimated for the PROGENy JAK-STAT z-score in the solid metastatic or intra-abdominal sample with the lowest JAK-STAT score at diagnosis with a multivariable Cox model adjusted for HRD status from SBS3 mutational signature, presence of pathogenic mutations in homologous recombination (HR) genes, high volume (> 1000 ml) of ascites at diagnosis and presence of macroscopic residual tumor after cytoreductive surgery as dichotomous variables, and, cancer dissemination score^15^ as a linear variable. For visualization with Kaplan-Meier survival estimator curves, the JAK-STAT z-scores were classified into 0-0.25, 0.25-0.75, and 0.75-1.00 quantiles in the subcohort of NACT-treated patients (*n* = 82 patients with bulk RNA-seq data). The separation of the curves was measured with a log-rank test.

### Functional cell lines experiments

#### Cell lines and reagents

Ovarian cancer cell line COV362 (Sigma-Aldrich) was cultured in Dulbecco’s Modified Eagle’s Medium (Gibco) supplemented with 10% (vol/vol) fetal bovine serum (FBS, Thermo Fisher Scientific) and 1% Penicillin-Streptomycin antibiotics (Thermo Fisher Scientific). Ovarian cancer cell line Kuramochi (JCRB Cell Bank) was cultured in RPMI 1640 (Gibco) supplemented with 10% FBS and 1% Penicillin-Streptomycin antibiotics. All cells were cultured at 37°C under 5% CO_2_ and were Mycoplasma-free. Recombinant Human Interferon alpha 2a (IFN-alpha) was purchased from R&D systems. Cisplatin was purchased from Sigma-Aldrich.

#### Cell viability assay

To assess the relative 50% inhibitory concentration (IC50) of COV362 or Kuramochi cells to cisplatin and IFN-alpha, cell viability assays were performed. Briefly, cells (5,000 cells/well) were seeded in 96-well plates and cultured overnight, then treated with either vehicle or different concentrations of cisplatin (0.5, 1, 2.5, 5, 10, 20, 30, 50μM) or IFN-alpha (0.01, 0.1, 1, 10, 100, 1000U/μl) for 72 h. For combination treatment, cells were treated with vehicle or 10U/μl of IFN-alpha combined with increasing concentration of cisplatin (0.5, 1, 2.5, 5, 10, 20, 30, 50μM) for 72h. Cell viability was assessed by measuring luminescence using CellTiter-Glo 2.0 Cell Viability Assay (Promega) in 2014 Envision multilabel reader (Perkin Elmer). Dose-response curves and relative IC50 were generated by GraphPad Prism 10. The IC50 curves of cisplatin for COV362 and Kuramochi cells are shown in Extended Data Fig. 3, and the IC50 curves of IFN-alpha for COV362 and Kuramochi cells are shown in Extended Data Fig. 3g.

#### Antibody-oligo conjugation

Oligonucleotides were conjugated to two antibodies against HGSOC cell surface proteins β2-Microglobulin (β2M) and CD298 (BioLegend) by iEDDA-click chemistry according to CITE-seq antibody-oligo conjugation protocol (https://cite-seq.com). Briefly, oligonucleotides were derivatized with TCO-PEG4-NHS (Click chemistry tools) in 10x borate buffered saline (BBS), and the TCO-labeled oligo was verified by Bioanalyzer Small RNA chip (Agilent). Antibodies were conjugated to mTz-PEG4-NHS (Click chemistry tools) at first in 1x BBS, then a 300μl reaction containing 300μg of mTz-PEG4-antibodies and 3nmol of TCO-PEG4-Oligo in 1x BBS was reacted at 4°C overnight. Excess mTz was quenched with 30μl of 10mM TCO-PEG4-glycine at room temperature for 10 min. Excess oligos were washed by Amicon Ultra-0.5 centrifugal filter unit with 50kDa MWCO membrane (Millipore) with 1x PBS. The conjugation efficiency was assessed by size shift of the conjugated antibodies on a 4-12% PAGE-gel (Thermo Fisher Scientific).

#### Single-cell RNA seq combined with cell hashing

This experiment was performed using Chromium Next GEM Single Cell 3’ Reagent Kits v3.1 (Dual Index) (10x Genomics, CG000317 Rev C). COV362 or Kuramochi cells were seeded in the 96-well plate, cultured overnight, and treated with cisplatin at different doses (10%, 50%, and 200% of IC50) and for different treatment times (24h, 12h). Subsequently, cells in different treatment conditions were labeled with distinct oligo-conjugated antibodies (Hashtags) against HGSC cell surface proteins β2-Microglobulin (β2M) and CD298. All cells were pooled and subjected to one 10x single-cell RNA seq run. The gene expression and hashtag libraries for NGS were constructed and sequenced according to the manufacturer’s protocol. Library quality was assessed on the Bioanalyser using the Agilent High Sensitivity DNA kit (Agilent). Libraries were sequenced on the Illumina NextSeq 2000 sequencing platform according to 10x manufacturer’s protocol. Raw base call files were demultiplexed into FASTQ files, aligned to the GRCh38 human reference genome, filtered, and processed to count barcodes and UMIs using the Cell Ranger software (v6.0.2, 10x Genomics). Cell demultiplexing was performed using the demultiplexing function HTODemux from the R package Seurat v4.2.0. The subsequent analysis used the single-cell filtered count matrix of all the genes.

#### Extraction of the cisplatin gene expression signature

In total, we had five replicated experiments for each cell line that included treatment with different concentrations of cisplatin and at different times. For each experiment independently, we calculated the effect size as Cohen’s d between cisplatin-treated cells (10%, 50%, and 200% concentration, 12h, and 24h) and control cells (control and vehicle) for each gene separately and obtained a vector containing the effect size between treated and control cells across all genes. This vector ranks the genes by how the cisplatin treatment affects them (from upregulated to downregulated), and we used it as a proxy for a ‘cisplatin gene expression signature.’ To evaluate the agreement between the 10 cisplatin gene expression signatures obtained, we calculated the Pearson correlation between all of them. To obtain the average cisplatin gene expression signature, we calculated the mean of effect sizes across the 10 experiments for each gene.

#### Classification of cells based on sensitivity to cisplatin

For each cell treated with cisplatin at a concentration of 200% during 24h, we calculated the cisplatin sensitivity score (CisSenScore) as the log ratio between the counts in the 1000 genes with higher effect size in the average cisplatin gene expression signature and the 10,000 genes with lower effect size (serves as background to control for the difference in total counts across cells). Based on this CisSenScore, we classified the treated cells (at concentration 200% for 24 h) into ‘more-sensitive’ (25% of cells with the highest CisSenScore) and ‘less-sensitive’ (25% of cells with the lowest CisSenScore).

#### Identifying the transcriptional programs associated with drug sensitivity

For each experiment, we applied a principal component analysis (PCA) to the expression values of the control cells. The expression values were centered separately for treated and control cells. As the PCs were calculated from gene expression values from the control experiments (i.e., without any perturbation), they reflect the background changes in gene expression. Then, we projected the expression values from the treated cells onto the PC space of the control cells. For the subsequent analyses, we used the first 10 PCs with the highest variance in PCA. Then, for each experiment, we checked which of the 10 PCs separate the ‘more-sensitive’ and the ‘less-sensitive’ cells in the projected data (i.e., values from the treated cells). Specifically, for each PC, we calculated the effect size (as Cohen’s *d*) between the PC activities of the cells previously classified as ‘more-sensitive’ versus the ‘less-sensitive.’ To account for the cell cycle effects, we also checked which of the 10 PCs separate the cell cycle phases (effect size between G1 vs. S, G1 vs G2M, and S vs G2M cells). For subsequent analyses, we selected 27 PCs with an absolute Cohen’s *d* greater than 0.5 between ‘more-sensitive’ and ‘less-sensitive’ cells, but that did not separate any of the cell cycle phases (Cohen’s *d* < 0.5 in all of the three cell cycle phases). Each of these selected PCs represents gene expression patterns or transcriptional programs (up-regulation and down-regulation of certain genes) of inter-individual variability in non-treated cells (only present in a subset of the non-treated cells). When these transcriptional programs are upregulated (or down-regulated) in the treated cells, they correlate with a higher (or lower) sensitivity to cisplatin. As a validation, we expected to identify the same transcriptional programs associated with drug sensitivity in the different replicated experiments. Therefore, we performed a hierarchical clustering of the 27 selected PCs that separated ‘more-sensitive’ and ‘less-sensitive’ cells. By visual inspection, we selected three clusters that capture three robust (found in several replicates) transcriptional programs associated with drug sensitivity. For each cluster, we calculated an average transcriptional program (mean value across all PCs in the cluster by gene). Then, we estimated the pathway activities induced by each of the three robust transcriptional programs (clusters) using GSEA^20^. For each cluster, we ranked the genes according to the median gene weight across the different gene expression signatures in the cluster and applied the GSEA function from the ClusterProfiler package for the hallmark gene sets of the Human MSigDB^21^ Collections.

## Data availability

All raw DNA sequencing data are submitted to the European Genome-phenome Archive (EGA) and will be publicly available under study accession number EGAS00001006775. Raw bulk RNA sequencing data are deposited in the EGA and are publicly available (EGAS00001004714). Quantified signals for t-CycIF data will be available in Synapse upon publication. Unpublished raw scRNA-seq data will be deposited in the EGA, and processed scRNA-seq data will be deposited in Gene Expression Omnibus (GEO) upon publication. Source data used in Fig. 3a-d and Extended Data Fig. 2a,c,d,e,f are provided in this paper. All other data supporting the results of this study are available from the corresponding author upon reasonable request.

## Code availability

The code related to the bulk RNA-seq and scRNA-seq analyses will be publicly available in the GitHub repository (https://github.com/XXX) upon publication. Specific code will be made available upon request to. Python and R packages used for this study are described in Methods. All packages are public and are freely available online.

## Acknowledgments

We thank the patients and their families. This project received funding from the European Union’s Horizon 2020 Research and Innovation Programme under grant agreement 965193 (DECIDER), the Research Council of Finland, the Sigrid Jusélius Foundation, the Cancer Foundation Finland, Cancerfonden (Sweden), and the Swedish Research Council. This study was co-funded by the European Union (ERC, SPACE 101076096). Views and opinions expressed are, however, those of the authors only and do not necessarily reflect those of the European Union or the European Research Council. Neither the European Union nor the granting authority can be held responsible for them. This programme has been supported with an educational grant via the Gilead Nordic Fellowship Programme. The authors wish to acknowledge the CSC-IT Center for Science (Finland) for computational resources. The Auria biobank (https://www.auria.fi/biopankki/) is acknowledged for delivering biobank samples and H&E images to our study. The authors also thank the core facility at NEO, BEA, Bioinformatics and Expression Analysis, which is supported by the board of research at the Karolinska Institutet and the research committee at the Karolinska Hospital. <u>BioRender.com</u> is acknowledged for providing illustration software using which Fig. 1a, Extended Data Fig.1, Extended Data Fig. 2a, and Extended Data Fig. 3a were made. We thank Dr. Kaisa Huhtinen for assisting in sample sequencing, Dr. Ann-Christin Ostwaldt for coordinating the DECIDER project, and Jenni Lahtinen for the management of laboratory operations.

## Author information

These authors contributed equally: Daria Afenteva, Rong Yu.

These authors jointly supervised the work: Sampsa Hautaniemi, Jussi Taipale.

D.A., R.Y., I.S., J.H., T.A.M., J.T., and S.Ha conceptualized the project. S.Hi and J.H. oversaw patient enrolment and coordinated the sample collection. A.R., V.-M.I., A.Vi, and J.H. reviewed clinical data. D.A., K.Z., and S.J. performed bulk RNA-seq sample processing and analysis. D.A., K.Z., E.P.E., M.M.F., and A.Vä performed scRNA-seq sample processing and analysis. K.L., G.Ma, Y.L., G.Mi, A.L., J.O., and T.M. performed WGS sample processing and analysis. I.-M.L., A.F. performed CycIF data processing and analysis. D.A. and T.A.M. performed and interpreted the omics computational analyses. R.Y., D.U., I.S., and J.T. designed and performed the cell lines experiments. M.S. analyzed and interpreted results from the cell line experiments. D.A., R.Y., M.S., I.S., and T.A.M. wrote the draft of the manuscript. D.A., R.Y., A.R., M.S., I.-M.L., K.L., G.Ma, G.Mi, I.S., J.H., T.A.M., J.T., and S.Ha contributed to the writing. All authors reviewed and approved the manuscript.

## Ethics declarations

All patients participating in the study gave their informed consent, and the study was approved by the Ethics Committee of the Hospital District of Southwest Finland (ETMK 145/1801/2015). The authors declare no competing interests.

## Supplementary information

**Supplementary Table 1.** Patient and tumor characteristics.

|  |  | chemo-refractory patients | chemo-sensitive patients | p-value *<br>(refractory vs. sensitive) | intermediate NACT-treated patients | PDS-treated patients |
| --- | --- | --- | --- | --- | --- | --- |
| **Diagnosis age** | median [range] | 70 [51-85] | 68 [38-86] | 0.97 | 68 [54-83] | 68.5 [39-88] |
| **Stage (FIGO2014)** | IIA-IIIA | 0 (0%) | 0 (0%) | 0.9 | 0 (0%) | 20 (14%) |
|  | IIIB | 0 (0%) | 0 (0%) |  | 1 (3%) | 8 (6%) |
|  | IIIC | 20 (65%) | 38 (61%) |  | 27 (69%) | 79 (56%) |
|  | IVA | 5 (16%) | 13 (21%) |  | 7 (18%) | 8 (6%) |
|  | IVB | 6 (19%) | 11 (18%) |  | 4 (10%) | 27 (19%) |
| **Dissemination score\*\*** | 1-5 | 0 (0%) | 3 (6%) | 0.026 | 1 (3%) | 37 (26%) |
|  | 6-10 | 7 (27%) | 12 (24%) |  | 8 (26%) | 61 (44%) |
|  | 11-15 | 15 (58%) | 34 (69%) |  | 18 (58%) | 38 (27%) |
|  | >15 | 4 (15%) | 0 (0%) |  | 4 (13%) | 4 (3%) |
|  | unknown | 5 (16%) | 13 (21%) |  | 8 (21%) | 2 (1%) |
| **Over 1000ml ascites at diagnosis** | No | 6 (19%) | 27 (44%) | 0.024 | 14 (36%) | 89 (64%) |
|  | Yes | 25 (81%) | 35 (56%) |  | 25 (64%) | 50 (36%) |
|  | unknown | 0 (0%) | 0 (0%) |  | 0 (0%) | 3 (2%) |
| **SBS3 mutational signature** | HRD | 11 (39%) | 28 (53%) | 0.35 | 10 (33%) | 56 (55%) |
|  | HRP | 17 (61%) | 25 (47%) |  | 20 (67%) | 45 (45%) |
|  | unknown | 3 (10%) | 9 (15%) |  | 9 (23%) | 41 (29%) |
| **ID6 mutation signature** | HRD | 7 (30%) | 25 (50%) | 0.14 | 4 (14%) | 55 (47%) |
|  | HRP | 16 (70%) | 25 (50%) |  | 24 (86%) | 62 (53%) |
|  | unknown | 8 (26%) | 12 (19%) |  | 11 (28%) | 25 (18%) |
| **HR gene mutation** | No | 20 (87%) | 39 (75%) | 0.36 | 26 (93%) | 86 (73%) |
|  | Yes | 3 (13%) | 13 (25%) |  | 2 (7%) | 32 (27%) |
|  | unknown | 8 (26%) | 10 (16%) |  | 11 (28%) | 24 (17%) |
| **Foldback inversions** | Not detected | 13 (59%) | 25 (49%) | 0.46 | 14 (50%) | 60 (51%) |
|  | Detected | 9 (41%) | 26 (51%) |  | 14 (50%) | 58 (49%) |
|  | unknown | 9 (29%) | 11 (18%) |  | 11 (28%) | 24 (17%) |
| **Any debulking surgery** | Yes | 11 (35%) | 60 (97%) | 1.3e-10 | 33 (85%) | 142 (100%) |
|  | No | 20 (65%) | 2 (3%) |  | 6 (15%) | 0 (0%) |
| **Residual tumor after debulking surgery** | No visible tumor | 4 (13%) | 18 (29%) | 1.7e-06 | 15 (38%) | 72 (51%) |
|  | 1 to 10mm | 6 (19%) | 35 (56%) |  | 14 (36%) | 46 (32%) |
|  | more than 10mm | 21 (68%) | 9 (15%) |  | 10 (26%) | 24 (17%) |

\* The refractory and reference patient groups were compared with the Mann-Whitney test for diagnosis age and the Fisher exact test for other characteristics.
\*\* Isoviita et al. 2019

**Supplementary Table 2.** Genomic characteristics of Kuramochi and COV362 cell lines.

| Cell line | TP53 | BRCA1 | BRCA2 | RB1 | MYC |
| --- | --- | --- | --- | --- | --- |
| **Kuramochi** | Mutation |  | Mutation |  | Amplification |
| **COV362** | Mutation | Mutation |  | Low mRNA expression | Amplification |

## Extended Data

**Supplementary Figure 1.**
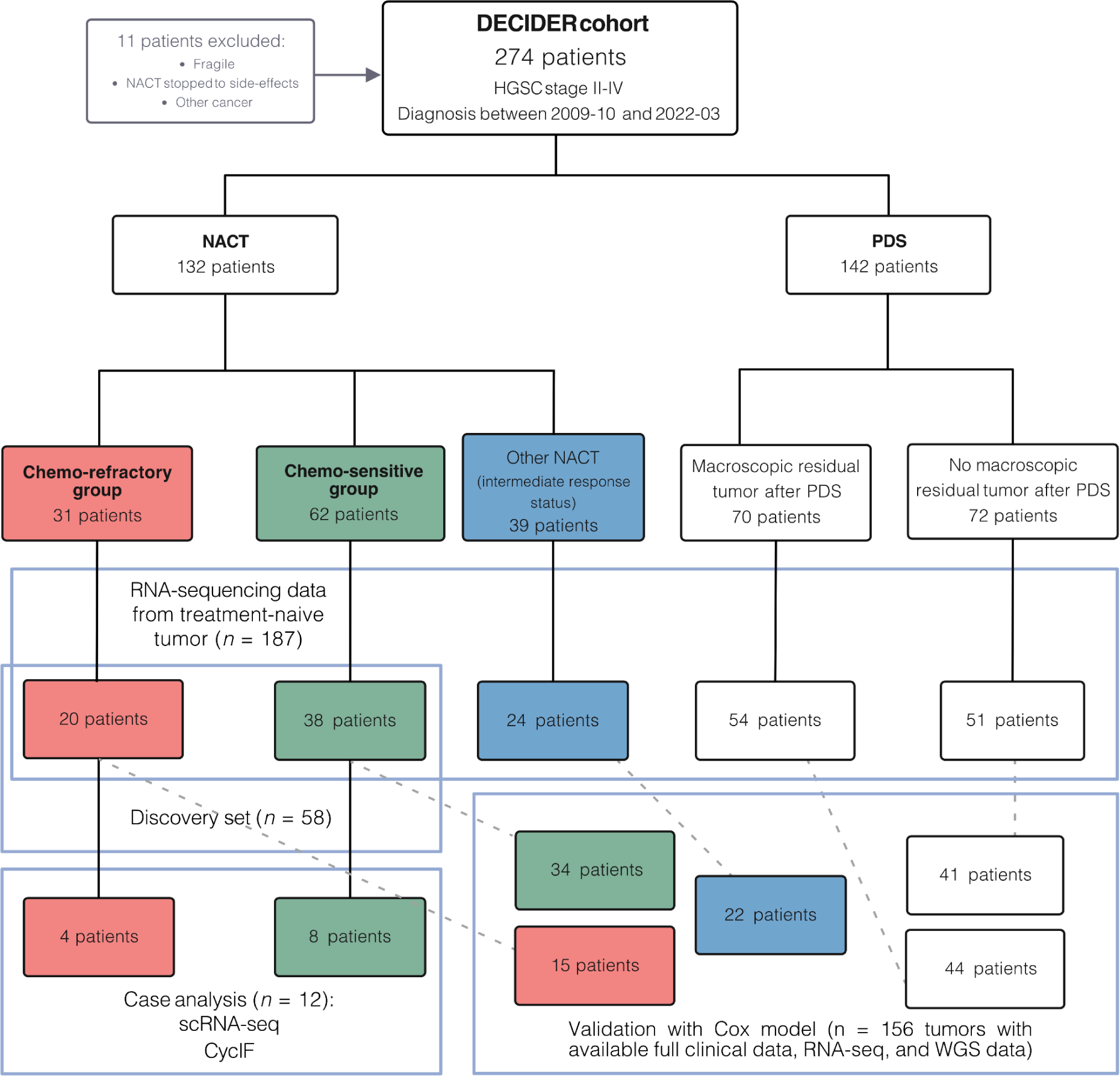
Flowchart of patient stratification from the DECIDER Clinical Trial. The chart illustrates the categorization of 274 high-grade serous ovarian cancer (HGSC) patients into 142 patients treated with primary debulking surgery (PDS) and 132 patients treated with neoadjuvant chemotherapy (NACT). The NACT group included chemo-refractory (*n* = 31), intermediate (*n* = 24), and chemo-sensitive (*n* = 62) patients based on the response to NACT and platinum-free interval (PFI), excluding 11 patients due to various reasons, such as inadequate treatment or other cancers (Methods). The discovery set comprised bulk RNA-sequencing data from 20 refractory and 38 sensitive patients, supplemented by single-cell RNA-seq (scRNA-seq) and cyclic immunofluorescence (CycIF) for a subset of patients. The set used for fitting the Cox proportional hazards model comprised additional NACT-treated patients with intermediate outcomes and patients who underwent PDS from the DECIDER cohort (*n* = 156).

**Supplementary Figure 2.**
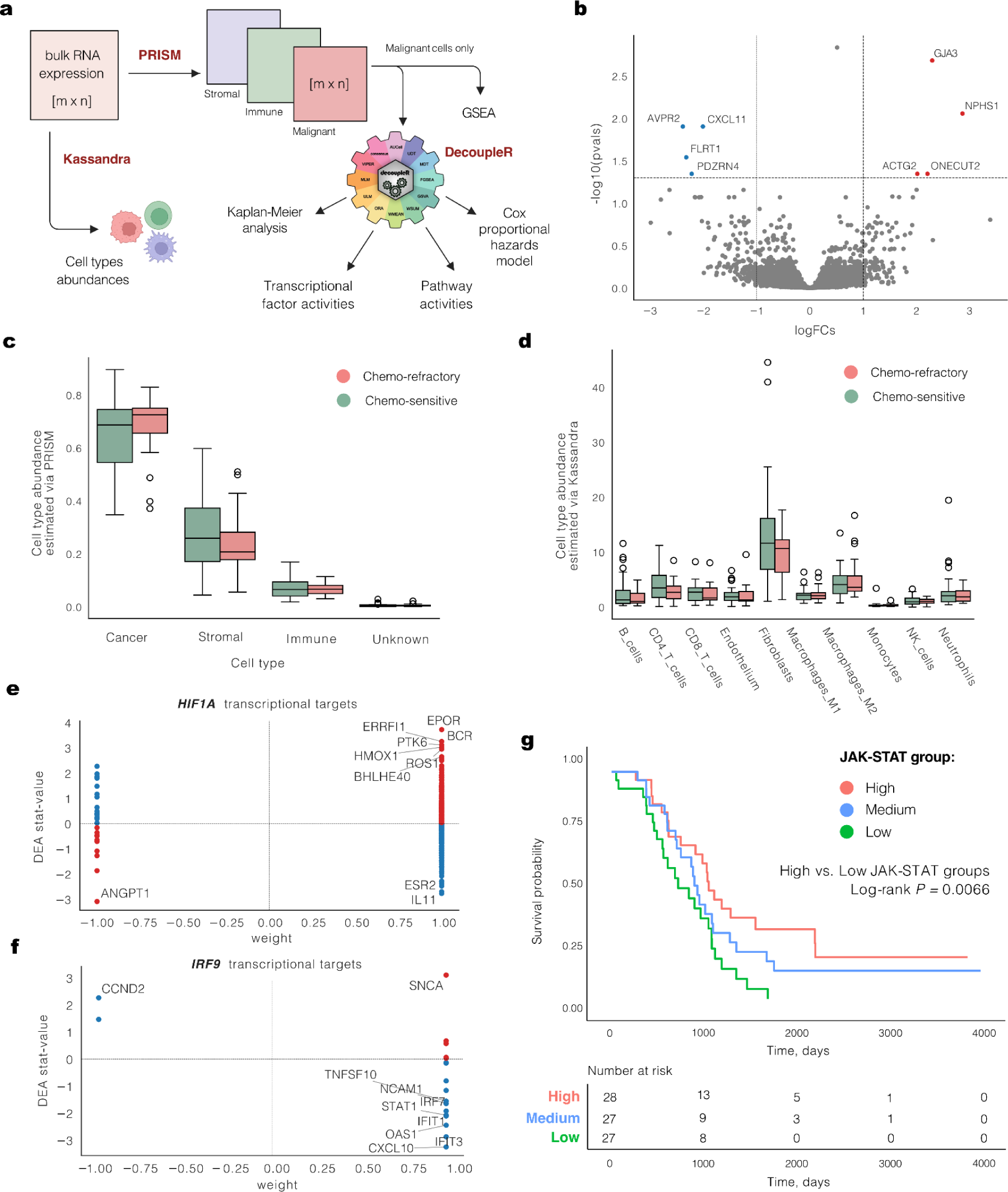
a, Workflow diagram of the bulk RNA expression analysis of samples from chemo-refractory (*n* = 20) and chemo-sensitive (*n* = 38) patients. b, The volcano plot of Differential Expression Analysis (DEA) results highlighting genes up-regulated (red) and down-regulated (blue) in the refractory tumors. c, A boxplot comparison of cell type abundances between chemo-refractory and chemo-sensitive patient samples according to PRISM. d, Boxplot comparison of cell type abundances between chemo-refractory and chemo-sensitive patient samples according to Kassandra. e, A scatter plot depicting the weight of the *HIF1A* transcriptional target genes (*x*-axis) and their stat-value derived from the DEA analysis (*y*-axis) of chemo-refractory versus chemo-sensitive tumors. f, Scatter plot depicting the weight of the *IRF9* transcriptional target genes (*x*-axis) and their stat-value derived from DEA analysis (*y*-axis) of chemo-refractory versus chemo-sensitive tumors. g, Kaplan-Meier survival curves stratified by JAK-STAT pathway activity levels into high (red), medium (blue), and low (green) in NACT-treated patients from the DECIDER cohort (*n* = 82 with available bulk RNA-seq data), with the number at risk table provided. Boxplots are presented as the range with the bounds of the box extending to the first and the third quantiles with a line at the median and whiskers extending to the farthest data point lying within 1.5x the interquartile range from the box.

**Supplementary Figure 3.**
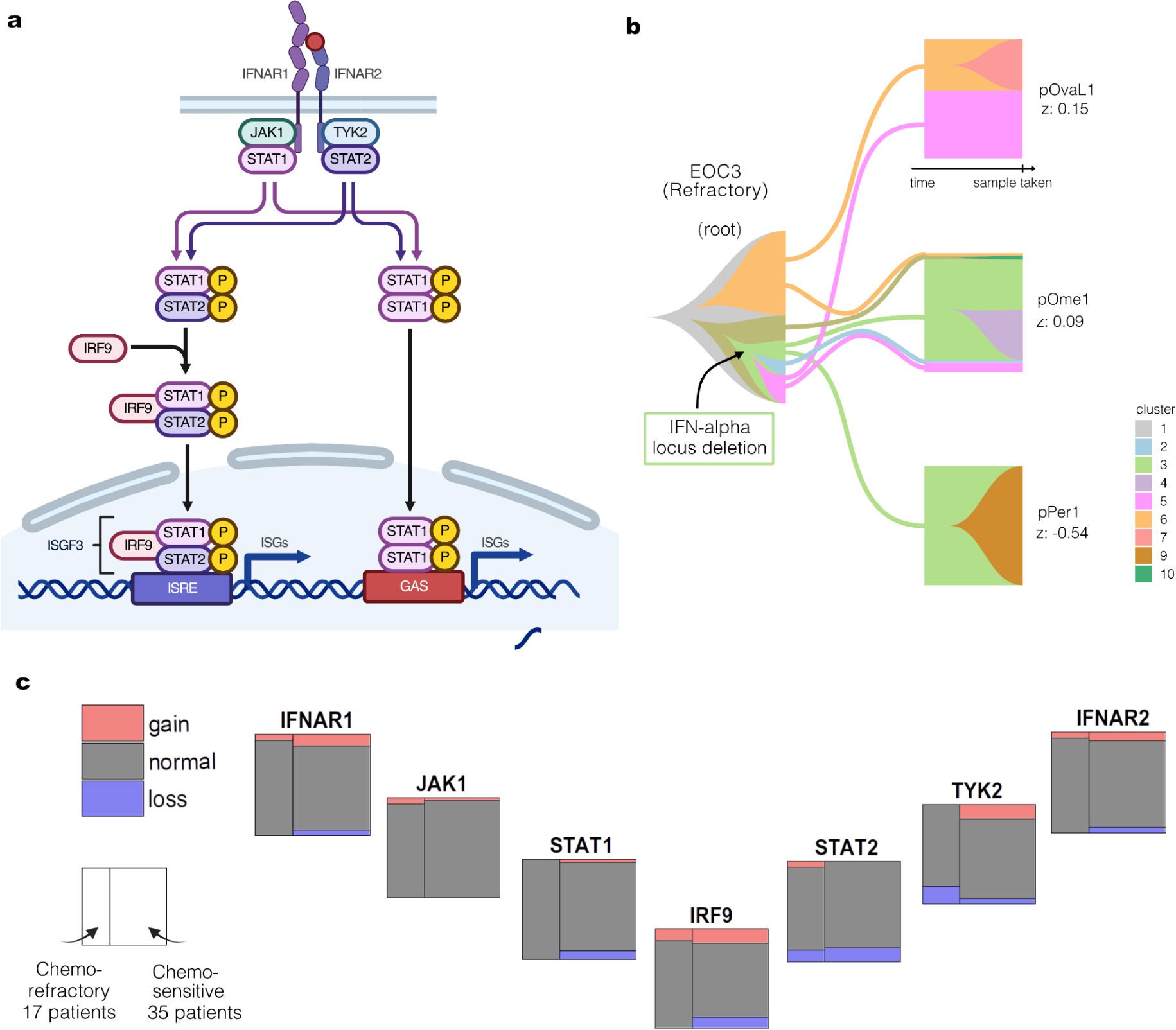
a, Schematic representation of the IFN-I signaling pathway, illustrating the cascade from IFNAR1/2 activation through JAK/STAT phosphorylation and ending with IRF9-mediated interferon-stimulated genes (ISGs) transcription. b, Jellyfish plot showing the clonal evolution across different samples in the chemo-refractory patient EOC3 harboring an IFN-alpha locus deletion, with colors indicating different subclones. The JAK-STAT z-scores per each sample are displayed as numbers. c, Mosaic plots showing genomic aberration analysis across the IFN-I signaling genes in chemo-refractory and chemo-sensitive HGSC patients. ISGF3, interferon-stimulated gene factor-3; ISRE, interferon-stimulated response element; GAS, gamma-activated sequence.

**Supplementary Figure 4.**
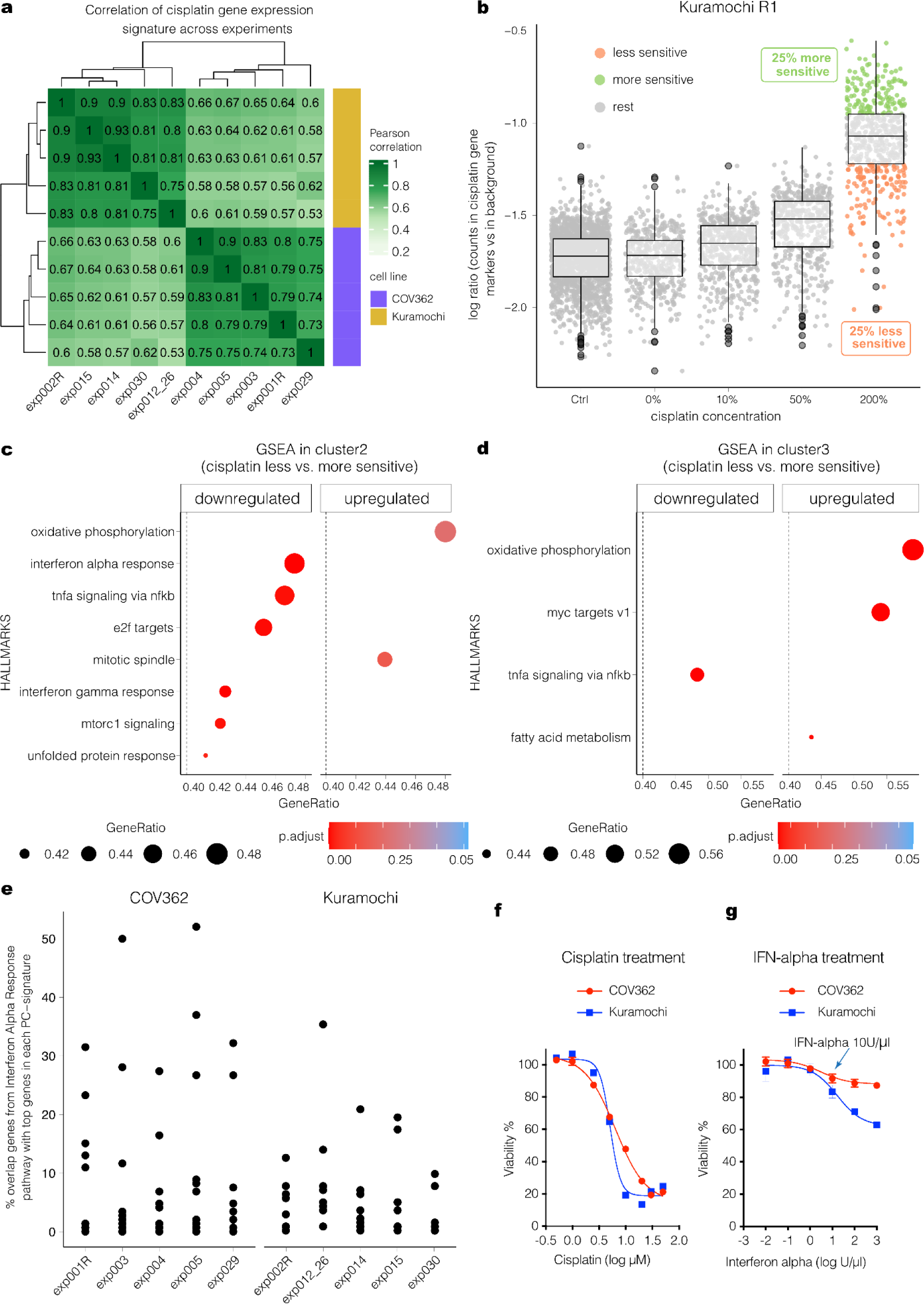
a, Hierarchical clustering of the correlation between the transcriptional signatures (gene weights) obtained from 10 experiments of both COV362 and Kuramochi. b, Based on the CisSenScore, the treated cells were separated into ‘more-sensitive’ (top 25%, green) and ‘less-sensitive’ (bottom 25%, orange) in one experiment of Kuramochi cells treated with cisplatin. c, d, GSEA scores for hallmark gene sets (with p.adjust < 0.05 and GeneRatio > 0.4) calculated based on the gene expression signatures in clusters 2 and 3 of the ‘less-sensitive’ cells compared with the ‘more-sensitive’ cells. e, Percentage of the 100 top genes in the Interferon Alpha Response pathway overlapping with top genes in each PC signature from the different experiments. Interferon Alpha Response variability signature was not recovered in Kuramochi cells due to their lack of baseline heterogeneity in the IFN-I response. f, Cell viability assay of cisplatin in ovarian cancer cells COV362 and Kuramochi. g, Cell viability assay of IFN-alpha in ovarian cancer cells COV362 and Kuramochi. Box plots are presented as the range (whiskers) with the bounds of the box extending to the first and the third quantiles with a line at the median and whiskers extending to the farthest data point lying within 1.5x the interquartile range from the box.

