## Supplementary material for "Multi-Omics Analysis Reveals the Attenuation of the Interferon Pathway as a Driver of Chemo-Refractory Ovarian Cancer": Source Data Table 1

| | JAK-STAT | Hypoxia |
|---|---|---|
| **EOC449** | 1,3042017 | -0,95886916 |
| **EOC1017** | -0,4859429 | 0,5777848 |
| **EOC888** | -1,8340892 | -0,2940742 |
| **EOC150** | -1,6831865 | 0,636615 |
| **EOC294** | -1,4784249 | 0,67190355 |
| **EOC26** | -1,9433738 | 0,23575453 |
| **EOC373** | 0,54216325 | 0,21353172 |
| **EOC197** | 0,9665245 | 0,42061654 |
| **EOC218** | 0,14253302 | -0,4990242 |
| **EOC1032** | -1,6375364 | 0,9016432 |
| **EOC691** | -0,5155427 | 0,9295721 |
| **EOC183** | 1,2314522 | -1,6501445 |
| **EOC49** | 0,8809856 | 0,17987369 |
| **EOC587** | 0,42454708 | 0,3610145 |
| **EOC991** | 0,9356111 | 0,026734952 |
| **EOC824** | -0,64525396 | -0,10988726 |
| **EOC649** | -0,4240256 | 1,6116253 |
| **EOC376** | -0,6224769 | -0,21542475 |
| **EOC933** | -2,0250268 | 1,0168471 |
| **EOC1129** | 0,20471378 | -0,31448627 |
| **EOC198** | -0,79612094 | 2,364416 |
| **EOC372** | 0,4360784 | -0,9013979 |
| **EOC423** | -1,4215782 | 1,0956926 |
| **EOC3** | -0,54237294 | 0,90836746 |
| **EOC87** | -0,36999878 | 1,1234412 |
| **EOC136** | -0,981168 | 1,2426744 |
| **EOC153** | -0,1023746 | 0,6358067 |
| **EOC658** | 0,8980579 | -1,7126849 |
| **EOC1127** | 0,7945114 | -0,22946668 |
| **EOC115** | -1,5256131 | 0,04431758 |
| **EOC227** | 0,052174743 | 1,5083162 |
| **EOC192** | 0,9590964 | -2,423278 |
| **EOC473** | 0,82201475 | 0,08910013 |
| **EOC1107** | -1,7012665 | -0,08448928 |
| **EOC257** | 0,14433576 | 1,0984977 |
| **EOC984** | -0,16284512 | 0,8179475 |
| **EOC175** | 0,9281363 | -0,4453707 |
| **EOC359** | 1,210351 | -0,15965717 |
| **EOC523** | -0,5394716 | 0,6588676 |
| **EOC612** | 1,2878681 | -1,8716364 |
| **EOC138** | -0,3527995 | 1,881788 |
| **EOC1038** | 1,1029049 | -0,8186063 |
| **EOC268** | -0,6879724 | 0,06409416 |
| **EOC864** | 1,2116295 | -0,29770076 |

| | | |
|---|---|---|
| **EOC551** | -0,85196114 | 0,20325679 |
| **EOC1023** | -0,9705672 | 0,3546334 |
| **EOC351** | 0,6290423 | -0,1235522 |
| **EOC60** | 0,5246969 | -0,175755 |
| **EOC323** | 0,4469761 | 0,719313 |
| **EOC1133** | -1,0460595 | 1,7375022 |
| **EOC768** | 1,655751 | -1,8118584 |
| **EOC1075** | 0,6910218 | -0,7693579 |
| **EOC1085** | -0,6973344 | 1,5355248 |
| **EOC868** | 1,071332 | -0,00431614 |
| **EOC958** | -0,86404675 | 0,9358499 |
| **EOC221** | 0,9128554 | 0,4920811 |
| **EOC295** | -1,4047347 | 0,7614329 |
| **EOC167** | -2,1976318 | -0,01386452 |
