## Supplementary material for "Multi-Omics Analysis Reveals the Attenuation of the Interferon Pathway as a Driver of Chemo-Refractory Ovarian Cancer": Source Data Table 2

| | EOC | Fibroblast | Immune | Unknown |
|---|---|---|---|---|
| **EOC1023** | 0,665346606 | 0,279912875 | 0,043360542 | 0,011379976 |
| **EOC868** | 0,702794112 | 0,193983849 | 0,100768463 | 0,002453576 |
| **EOC551** | 0,714216574 | 0,179822177 | 0,102315591 | 0,003645658 |
| **EOC60** | 0,627727343 | 0,286339732 | 0,081069411 | 0,004863514 |
| **EOC1133** | 0,635217316 | 0,266280581 | 0,09596001 | 0,002542093 |
| **EOC26** | 0,771251403 | 0,15588229 | 0,069635663 | 0,003230644 |
| **EOC218** | 0,391728818 | 0,410220822 | 0,169470296 | 0,028580064 |
| **EOC1032** | 0,746058713 | 0,174203384 | 0,071816336 | 0,007921568 |
| **EOC167** | 0,398099886 | 0,498266656 | 0,081503621 | 0,022129836 |
| **EOC183** | 0,544993618 | 0,352937057 | 0,095251942 | 0,006817383 |
| **EOC376** | 0,757439762 | 0,197000144 | 0,04256626 | 0,002993834 |
| **EOC1129** | 0,517661379 | 0,40156468 | 0,065545726 | 0,015228215 |
| **EOC372** | 0,546131098 | 0,37979256 | 0,068916967 | 0,005159375 |
| **EOC423** | 0,696892137 | 0,236684732 | 0,064017716 | 0,002405414 |
| **EOC87** | 0,487428355 | 0,428313877 | 0,080334454 | 0,003923314 |
| **EOC295** | 0,659362254 | 0,300485509 | 0,035724263 | 0,004427974 |
| **EOC587** | 0,37095791 | 0,510220097 | 0,106896159 | 0,011925834 |
| **EOC649** | 0,726065188 | 0,166975886 | 0,097193622 | 0,009765304 |
| **EOC933** | 0,872410037 | 0,071725555 | 0,054605451 | 0,001258957 |
| **EOC691** | 0,797677387 | 0,13011701 | 0,071814043 | 0,00039156 |
| **EOC49** | 0,409675794 | 0,539489218 | 0,037809962 | 0,013025026 |
| **EOC3** | 0,583236848 | 0,377715205 | 0,033556069 | 0,005491878 |
| **EOC136** | 0,64008884 | 0,338893044 | 0,01828589 | 0,002732226 |
| **EOC153** | 0,539195313 | 0,420987345 | 0,033707388 | 0,006109954 |
| **EOC1127** | 0,492342351 | 0,390999865 | 0,112733972 | 0,003923812 |
| **EOC115** | 0,726990941 | 0,238977895 | 0,032101855 | 0,001929309 |
| **EOC192** | 0,780229015 | 0,151517077 | 0,065582393 | 0,002671515 |
| **EOC197** | 0,473693386 | 0,450776835 | 0,067421257 | 0,008108522 |
| **EOC991** | 0,629702166 | 0,241473372 | 0,125066036 | 0,003758426 |
| **EOC658** | 0,34780381 | 0,597752345 | 0,049730284 | 0,00471356 |
| **EOC227** | 0,518511372 | 0,380934233 | 0,08538347 | 0,015170925 |
| **EOC473** | 0,551455684 | 0,346365291 | 0,094919157 | 0,007259869 |
| **EOC1107** | 0,69362151 | 0,27466649 | 0,028219666 | 0,003492334 |
| **EOC1075** | 0,735377187 | 0,104277315 | 0,158066525 | 0,002278973 |
| **EOC373** | 0,76396637 | 0,19474307 | 0,037226544 | 0,004064017 |
| **EOC198** | 0,672839031 | 0,252789206 | 0,062606293 | 0,011765471 |
| **EOC175** | 0,631056391 | 0,272541148 | 0,092520798 | 0,003881663 |
| **EOC612** | 0,798839778 | 0,043982837 | 0,154929825 | 0,00224756 |
| **EOC449** | 0,896013168 | 0,063197008 | 0,040214538 | 0,000575287 |
| **EOC1017** | 0,506643067 | 0,44858336 | 0,039397992 | 0,005375581 |
| **EOC888** | 0,823712974 | 0,135798309 | 0,037365235 | 0,003123482 |
| **EOC294** | 0,867497938 | 0,110708573 | 0,020719183 | 0,001074306 |
| **EOC257** | 0,680833445 | 0,259150527 | 0,05579299 | 0,004223038 |
| **EOC984** | 0,723908124 | 0,215031651 | 0,058164736 | 0,00289549 |

| | | | | |
|---|---|---|---|---|
| **EOC359** | 0,794463599 | 0,144922788 | 0,057981127 | 0,002632486 |
| **EOC523** | 0,829563595 | 0,055803383 | 0,114103147 | 0,000529875 |
| **EOC138** | 0,732738433 | 0,198929768 | 0,066455104 | 0,001876695 |
| **EOC1038** | 0,628050827 | 0,290932495 | 0,077276412 | 0,003740265 |
| **EOC268** | 0,745912621 | 0,189517463 | 0,060891164 | 0,003678751 |
| **EOC864** | 0,703079306 | 0,239074623 | 0,054567676 | 0,003278394 |
| **EOC351** | 0,692196978 | 0,259137171 | 0,045711811 | 0,00295404 |
| **EOC323** | 0,726462996 | 0,195507761 | 0,073493496 | 0,004535747 |
| **EOC1085** | 0,738324867 | 0,145707725 | 0,112692488 | 0,00327492 |
| **EOC958** | 0,767733815 | 0,200227305 | 0,030233209 | 0,001805671 |
| **EOC150** | 0,844700734 | 0,114351037 | 0,038715537 | 0,002232692 |
| **EOC824** | 0,739475598 | 0,181751144 | 0,076729901 | 0,002043358 |
| **EOC768** | 0,748189685 | 0,182928391 | 0,064589715 | 0,004292209 |
| **EOC221** | 0,693086858 | 0,252784811 | 0,052030479 | 0,002097853 |
