## Supplementary material for "Multi-Omics Analysis Reveals the Attenuation of the Interferon Pathway as a Driver of Chemo-Refractory Ovarian Cancer": Source Data Table 3

| | B_cells | CD4_T_cells | CD8_T_cells | Endothelium | Fibroblasts | Macrophages_M1 | Macrophages_M2 | Monocytes | NK_cells | Neutrophils | Other |
|---|---|---|---|---|---|---|---|---|---|---|---|
| **EOC1023** | 0,88 | 3,24 | 1,5 | 2,86 | 11,09 | 2,55 | 3,65 | 0,2 | 0,91 | 2,06 | 71,04 |
| **EOC868** | 0,36 | 4,78 | 2,95 | 1,36 | 8,15 | 2,51 | 2,8 | 0,4 | 1,72 | 2,26 | 72,7 |
| **EOC551** | 4,28 | 3,62 | 2,68 | 0,93 | 10,48 | 2,28 | 3,19 | 0,24 | 1,5 | 3,76 | 67,04 |
| **EOC60** | 2,68 | 5,63 | 8,07 | 1,97 | 5,65 | 5,42 | 8,71 | 0,66 | 1,98 | 4,71 | 54,52 |
| **EOC1133** | 0,3 | 2,31 | 3,27 | 0,47 | 8,97 | 3,31 | 4,28 | 0,52 | 0,69 | 2,81 | 73,06 |
| **EOC26** | 0,6 | 0,15 | 0,36 | 0,93 | 10,84 | 0,78 | 1,82 | 0,24 | 0,05 | 1,36 | 82,87 |
| **EOC218** | 3,46 | 6,25 | 4,78 | 4,85 | 14,88 | 3,88 | 13,48 | 3,42 | 1,25 | 19,47 | 24,28 |
| **EOC1032** | 1 | 0,89 | 1,36 | 4,04 | 12,11 | 1,57 | 4,02 | 0,37 | 1,35 | 1,77 | 71,53 |
| **EOC167** | 0,73 | 4,49 | 4,55 | 9,56 | 15,97 | 4,25 | 12,05 | 0,49 | 1,39 | 2,08 | 44,45 |
| **EOC183** | 6,38 | 10,03 | 8,26 | 2,84 | 13,86 | 1,52 | 6,64 | 0,61 | 1,15 | 2,65 | 46,06 |
| **EOC376** | 1,14 | 1,78 | 1,07 | 1,27 | 11,19 | 1,56 | 1,98 | 0,34 | 0,56 | 1,29 | 77,83 |
| **EOC1129** | 7,1 | 6,66 | 2,14 | 6,66 | 5,22 | 1,48 | 3,13 | 0,51 | 1,16 | 1,3 | 64,63 |
| **EOC372** | 3,42 | 11,23 | 4,33 | 2,7 | 23,34 | 2,91 | 5,59 | 0,27 | 1,76 | 2 | 42,46 |
| **EOC423** | 0,82 | 1,73 | 1,13 | 1,88 | 16,58 | 1,26 | 1,74 | 0,46 | 0,51 | 0,72 | 73,17 |
| **EOC87** | 0,54 | 2,99 | 3,37 | 1,25 | 12,03 | 2,19 | 4,59 | 0,26 | 1,31 | 1,99 | 69,47 |
| **EOC295** | 0,65 | 1,05 | 0,92 | 2,59 | 12,72 | 1,33 | 3,08 | 0,35 | 0,4 | 0,9 | 76,01 |
| **EOC587** | 1,29 | 8,49 | 2,76 | 3,67 | 17,71 | 6,28 | 16,69 | 0,71 | 0,68 | 4,52 | 37,2 |
| **EOC649** | 11,59 | 6,06 | 3,85 | 3,38 | 6,64 | 1,14 | 3,13 | 0,19 | 1,43 | 2,64 | 59,96 |
| **EOC933** | 0,31 | 0,1 | 0,37 | 0,89 | 2,43 | 1,12 | 2,22 | 0,26 | 0,08 | 0,48 | 91,75 |
| **EOC691** | 0,69 | 2,42 | 1,85 | 0,13 | 7,53 | 2,03 | 5,83 | 0,3 | 3,29 | 2,52 | 73,41 |
| **EOC49** | 1,28 | 4,99 | 3,21 | 4,96 | 44,57 | 2,15 | 5,76 | 0,26 | 0,9 | 1,29 | 30,63 |
| **EOC3** | 0,97 | 2,81 | 1,16 | 2,81 | 15,89 | 1,81 | 4,86 | 0,15 | 0,73 | 1,68 | 67,14 |
| **EOC136** | 0,44 | 0,61 | 0,57 | 2,32 | 18,01 | 1,25 | 2,06 | 0,13 | 0,22 | 0,42 | 73,96 |
| **EOC153** | 5,6 | 4,49 | 1,99 | 3,12 | 21,49 | 1,73 | 4,72 | 0,13 | 0,57 | 2,09 | 54,06 |
| **EOC1127** | 2,87 | 3,16 | 3,44 | 1,71 | 18,76 | 2,46 | 5,21 | 0,11 | 0,88 | 2,81 | 58,59 |
| **EOC115** | 2,04 | 2,09 | 3,81 | 1,25 | 11,4 | 2,25 | 2 | 0,2 | 1,99 | 1,53 | 71,44 |
| **EOC192** | 0,99 | 2,22 | 1,3 | 1,84 | 6,12 | 1,89 | 2,34 | 0,38 | 0,98 | 1,65 | 80,29 |
| **EOC197** | 1,68 | 5,71 | 3,77 | 3,54 | 21,37 | 3,38 | 5,64 | 0,09 | 2,81 | 2,21 | 49,81 |
| **EOC991** | 2,41 | 1,94 | 2,68 | 1 | 12,75 | 6,37 | 5,9 | 0,22 | 0,79 | 4,86 | 61,06 |
| **EOC658** | 1,57 | 3,98 | 3,12 | 3,32 | 40,98 | 2,5 | 3,92 | 0,14 | 0,79 | 3,22 | 36,47 |
| **EOC227** | 8,67 | 5,83 | 3,41 | 5,25 | 14,85 | 1,43 | 8,51 | 0,3 | 2,13 | 7,21 | 42,41 |
| **EOC473** | 3,09 | 7,32 | 2,83 | 1,94 | 12,06 | 2,42 | 7,66 | 0,27 | 2,43 | 3,76 | 56,22 |
| **EOC1107** | 0,59 | 1,32 | 1,7 | 2,09 | 13,17 | 0,97 | 1,84 | 0,3 | 0,76 | 0,83 | 76,42 |
| **EOC1075** | 2,58 | 4,44 | 2,86 | 0,35 | 6,34 | 3,63 | 4,78 | 0,15 | 1,94 | 2,7 | 70,23 |
| **EOC373** | 2,95 | 1,77 | 1,28 | 2,58 | 10,35 | 1,08 | 2,13 | 0,3 | 1,64 | 2,06 | 73,86 |
| **EOC198** | 3,67 | 1,11 | 0,96 | 3,02 | 14,74 | 3,57 | 6,78 | 1,22 | 0,67 | 3,51 | 60,74 |
| **EOC175** | 9,06 | 8,62 | 4,77 | 1,49 | 14,33 | 1,65 | 3,18 | 0,22 | 1,64 | 1,1 | 53,94 |
| **EOC612** | 0,27 | 3,8 | 3,56 | 0,13 | 1,05 | 2,4 | 3,22 | 0,34 | 2,65 | 0,56 | 82,03 |
| **EOC449** | 0,65 | 1,11 | 0,72 | 0,16 | 2,08 | 1,09 | 1,57 | 0,22 | 0,46 | 0,52 | 91,42 |
| **EOC1017** | 0,7 | 5,9 | 1,55 | 2,53 | 25,54 | 2,53 | 5,65 | 0,08 | 0,72 | 1,18 | 53,59 |
| **EOC888** | 0,43 | 0,27 | 1,77 | 2,05 | 4,83 | 0,7 | 1,6 | 0,41 | 0,56 | 8,39 | 78,98 |
| **EOC294** | 0,91 | 0,25 | 0,36 | 0,36 | 5,67 | 0,82 | 0,78 | 0,08 | 0,06 | 1,62 | 89,09 |
| **EOC257** | 1,63 | 2,81 | 5,09 | 1,86 | 17,38 | 2,23 | 6,24 | 0,28 | 0,77 | 3 | 58,71 |
| **EOC984** | 0,65 | 1,59 | 1,85 | 1,15 | 10,18 | 1,41 | 3,56 | 0,45 | 0,53 | 1,1 | 77,52 |
| **EOC359** | 2,24 | 2,76 | 1,32 | 0,85 | 4,46 | 2 | 3,1 | 0,22 | 1,37 | 0,75 | 80,94 |
| **EOC523** | 0,25 | 1,66 | 1,51 | 0,22 | 1,37 | 2,68 | 11,57 | 0,33 | 0,78 | 1,04 | 78,58 |
| **EOC138** | 0,41 | 0,35 | 0,56 | 0,58 | 10,69 | 3,19 | 5,95 | 0,26 | 0,53 | 5,14 | 72,35 |
| **EOC1038** | 2,5 | 6,94 | 3,67 | 1,59 | 13,41 | 4,59 | 8,25 | 0,13 | 1,32 | 2,63 | 54,97 |
| **EOC268** | 4,94 | 5,24 | 3,22 | 1,67 | 7,39 | 2,59 | 3,19 | 0,23 | 1,06 | 2,8 | 67,67 |
| **EOC864** | 0,87 | 2,03 | 1,18 | 1,89 | 8,72 | 2,37 | 4,88 | 0,26 | 1,19 | 8,09 | 68,51 |
| **EOC351** | 1,38 | 4,44 | 3,91 | 1,46 | 10,12 | 2,25 | 3,55 | 0,46 | 1,62 | 0,69 | 70,14 |
| **EOC323** | 3,14 | 4,78 | 2,81 | 2,23 | 9,87 | 3,98 | 5,21 | 0,4 | 1,04 | 1,47 | 65,07 |
| **EOC1085** | 0,82 | 1,44 | 0,96 | 1,18 | 6,51 | 1,41 | 2,3 | 0,56 | 0,8 | 0,96 | 83,05 |
| **EOC958** | 0,7 | 1,35 | 1,34 | 1,25 | 12,73 | 1,3 | 1,83 | 0,23 | 0,36 | 0,7 | 78,22 |
| **EOC150** | 0,28 | 0,14 | 0,3 | 1,21 | 4,49 | 0,82 | 1,89 | 0,17 | 0,03 | 0,45 | 90,21 |
| **EOC824** | 1,07 | 2,62 | 3,89 | 1 | 5,05 | 1,89 | 3,27 | 0,11 | 1,02 | 0,88 | 79,18 |
| **EOC768** | 2,69 | 3,22 | 2,01 | 2,04 | 9,85 | 2,15 | 2,9 | 0,17 | 2,01 | 0,79 | 72,17 |
| **EOC221** | 2,46 | 5,21 | 4,44 | 1,41 | 5,91 | 1,55 | 5,28 | 0,27 | 1,55 | 4,95 | 66,97 |
