## Supplementary Table 3 for "Multi-Omics Analysis Reveals the Attenuation of the Interferon Pathway as a Driver of Chemo-Refractory Ovarian Cancer"

| | | |
|---|---|---|
| Patient | Patient code | EOCX |
| Stratum | Status of the patient in the study | Refractory/Responsive |
| RNA sample in discovery set | Bulk RNA-seq sample that was taken into the discovery analysis (pathways and transcriptional factors activities) | EOCX_tissue |
| RNA sample with lowest JAK-STAT | Bulk RNA-seq sample that was used for Cox proportional hazards model and Kaplan-Meier estimator | EOCX_tissue |
| WGS_data | Whether WGS data from diagnostic sample was available for this patient | Yes/NA |
| scRNA_data | Whether scRNA-seq data from diagnostic sample was available for this patient | Yes/NA |
| CycIF_data | Whether CycIF data from diagnostic sample was available for this patient | Yes/NA |
| clinical data | Whether any clinical data was available for the patient. FIGO2014 stage, dissemination score, ascites at diagnosis, residual tumor after cytoreductive surgery. | Yes/NA |

| Patient | Stratum | RNA sample in discovery set | RNA sample with lowest JAK-STAT | WGS_data | scRNA_data | CycIF_data | clinical data |
|---|---|---|---|---|---|---|---|
| EOC868 | Sensitive | EOC868_Per | EOC868_Per | Yes | | | Yes |
| EOC1133 | Sensitive | EOC1133_Per | EOC1133_Per | Yes | | | Yes |
| EOC218 | Sensitive | EOC218_Ome | EOC218_Ome | Yes | | | Yes |
| EOC183 | Sensitive | EOC183_Ome | EOC183_Per | Yes | | | Yes |
| EOC376 | Sensitive | EOC376_Ome | EOC376_Ome | Yes | | | Yes |
| EOC1129 | Sensitive | EOC1129_Ome | EOC1129_Ome | Yes | | | Yes |
| EOC372 | Sensitive | EOC372_Per | EOC372_Per | Yes | Yes | | Yes |
| EOC423 | Sensitive | EOC423_Ome | EOC423_Ome | Yes | | | Yes |
| EOC295 | Sensitive | EOC295_Per | EOC295_Per | Yes | | | Yes |
| EOC649 | Sensitive | EOC649_Ome | EOC649_Ome | Yes | Yes | | Yes |
| EOC933 | Sensitive | EOC933_Ome | EOC933_Ova | Yes | | Yes | Yes |
| EOC691 | Sensitive | EOC691_Per | EOC691_Per | Yes | | | Yes |
| EOC49 | Sensitive | EOC49_Ome | EOC49_Ome | Yes | | | Yes |
| EOC136 | Sensitive | EOC136_Per | EOC136_Per | Yes | Yes | | Yes |
| EOC153 | Sensitive | EOC153_Ome | EOC153_Per | Yes | Yes | | Yes |
| EOC1127 | Sensitive | EOC1127_Ome | EOC1127_Ome | Yes | Yes | | Yes |
| EOC192 | Sensitive | EOC192_Ome | EOC192_Ova | Yes | | | Yes |
| EOC197 | Sensitive | EOC197_Per | EOC197_Tub | Yes | | | Yes |
| EOC991 | Sensitive | EOC991_Ome | EOC991_Ome | Yes | | | Yes |
| EOC658 | Sensitive | EOC658_Per | EOC658_Per | Yes | | | Yes |
| EOC227 | Sensitive | EOC227_Ome | EOC227_Ome | Yes | Yes | | Yes |
| EOC473 | Sensitive | EOC473_Ome | EOC473_Per | Yes | | | Yes |
| EOC1107 | Sensitive | EOC1107_Ome | EOC1107_Ome | Yes | | | Yes |
| EOC1075 | Sensitive | EOC1075_Tub | EOC1075_Tub | Yes | | | Yes |
| EOC175 | Sensitive | EOC175_Ome | EOC175_Ome | Yes | | | Yes |
| EOC612 | Sensitive | EOC612_Ome | EOC612_Ome | Yes | | | Yes |
| EOC449 | Sensitive | EOC449_Per | EOC449_Per | Yes | | | Yes |
| EOC1017 | Sensitive | EOC1017_Ome | EOC1017_Ome | Yes | | | Yes |
| EOC888 | Sensitive | EOC888_Ome | EOC888_Ome | Yes | | | Yes |
| EOC294 | Sensitive | EOC294_Per | EOC294_Per | Yes | | | Yes |
| EOC257 | Sensitive | EOC257_Per | EOC257_Per | Yes | | | Yes |
| EOC138 | Sensitive | EOC138_Per | EOC138_Per | Yes | | | Yes |
| EOC1038 | Sensitive | EOC1038_Tub | EOC1038_Tub | Yes | | | Yes |
| EOC864 | Sensitive | EOC864_Oth | EOC864_Oth | Yes | | | Yes |
| EOC351 | Sensitive | EOC351_Per | EOC351_Per | Yes | | | Yes |
| EOC323 | Sensitive | EOC323_Ova | EOC323_Oth | Yes | | | Yes |
| EOC150 | Sensitive | EOC150_Tub | EOC150_Tub | Yes | | | Yes |
| EOC768 | Sensitive | EOC768_Per | EOC768_Per | Yes | | | Yes |
| EOC940 | Sensitive | | | Yes | | Yes | Yes |
| EOC52 | Sensitive | | | Yes | | | Yes |
| EOC1122 | Sensitive | | | Yes | | | Yes |
| EOC955 | Sensitive | | | Yes | | | Yes |
| EOC599 | Sensitive | | | Yes | | | Yes |
| EOC389 | Sensitive | | | Yes | | | Yes |
| EOC6 | Sensitive | | | Yes | | | Yes |
| EOC928 | Sensitive | | | Yes | | | Yes |
| EOC15 | Sensitive | | | Yes | | | Yes |
| EOC600 | Sensitive | | | Yes | | | Yes |
| EOC914 | Sensitive | | | Yes | | | Yes |
| EOC881 | Sensitive | | | Yes | | | Yes |

| | | | | | | | |
|---|---|---|---|---|---|---|---|
| EOC286 | Sensitive | | | Yes | | | Yes |
| EOC454 | Sensitive | | | Yes | | | Yes |
| EOC806 | Sensitive | | | Yes | | | Yes |
| EOC568 | Sensitive | | | Yes | | | Yes |
| EOC1037 | Sensitive | | | Yes | | | Yes |
| EOC402 | Sensitive | | | | | | Yes |
| EOC96 | Sensitive | | | | | | Yes |
| EOC859 | Sensitive | | | | | | Yes |
| EOC698 | Sensitive | | | | | | Yes |
| EOC966 | Sensitive | | | | | | Yes |
| EOC993 | Sensitive | | | | | | Yes |
| EOC104 | Sensitive | | | | | | Yes |
| EOC1023 | Refractory | EOC1023_Ome | EOC1023_Ome | Yes | | | Yes |
| EOC551 | Refractory | EOC551_Ova | EOC551_Ova | Yes | | | Yes |
| EOC60 | Refractory | EOC60_Per | EOC60_Ome | Yes | | | Yes |
| EOC26 | Refractory | EOC26_Ome | EOC26_Ome | Yes | | | Yes |
| EOC1032 | Refractory | EOC1032_Per | EOC1032_Per | Yes | | | Yes |
| EOC167 | Refractory | EOC167_Ome | EOC167_Tum | Yes | | | Yes |
| EOC87 | Refractory | EOC87_Ome | EOC87_Ome | Yes | Yes | Yes | Yes |
| EOC587 | Refractory | EOC587_Adn | EOC587_Adn | Yes | | | Yes |
| EOC3 | Refractory | EOC3_Ome | EOC3_Per | Yes | Yes | Yes | Yes |
| EOC115 | Refractory | EOC115_Per | EOC115_Per | Yes | Yes | | Yes |
| EOC373 | Refractory | EOC373_Adn | EOC373_Adn | Yes | | | Yes |
| EOC198 | Refractory | EOC198_Per | EOC198_Per | Yes | | | Yes |
| EOC984 | Refractory | EOC984_Ome | EOC984_Per | Yes | | | Yes |
| EOC359 | Refractory | EOC359_Ome | EOC359_Per | Yes | | | Yes |
| EOC523 | Refractory | EOC523_Ome | EOC523_Ova | Yes | | | Yes |
| EOC268 | Refractory | EOC268_Per | EOC268_Per | Yes | | | Yes |
| EOC1085 | Refractory | EOC1085_Per | EOC1085_Per | Yes | | | Yes |
| EOC958 | Refractory | EOC958_Per | EOC958_Per | Yes | | | Yes |
| EOC824 | Refractory | EOC824_Per | EOC824_Per | Yes | | | Yes |
| EOC221 | Refractory | EOC221_Per | EOC221_Per | Yes | | | Yes |
| EOC204 | Refractory | | | Yes | Yes | | Yes |
| EOC895 | Refractory | | | Yes | | | Yes |
| EOC670 | Refractory | | | Yes | | | Yes |
| EOC889 | Refractory | | | Yes | | | Yes |
| EOC1007 | Refractory | | | Yes | | | Yes |
| EOC554 | Refractory | | | Yes | | | Yes |
| EOC9 | Refractory | | | Yes | | | Yes |
| EOC386 | Refractory | | | Yes | | | Yes |
| EOC1010 | Refractory | | | Yes | | | Yes |
| EOC46 | Refractory | | | | | | Yes |
| EOC194 | Refractory | | | | | | Yes |
